# A pair of esterases from a commensal gut bacterium remove acetylations from all positions on complex β-mannans

**DOI:** 10.1101/788067

**Authors:** Leszek Michalak, Sabina Leanti La Rosa, Shaun Allan Leivers, Lars Jordhøy Lindstad, Åsmund Røhr Kjendseth, Finn Lillelund Aachmann, Bjørge Westereng

## Abstract

β-Mannans and xylans are important components of the plant cell wall and they are acetylated to be protected from degradation by glycoside hydrolases. β-Mannans are widely present in human and animal diets as fiber from leguminous plants and as thickeners and stabilizers in processed foods. There are many fully characterized acetylxylan esterases (AcXEs), however, the enzymes deacetylating mannans are less understood. Here we present two carbohydrate esterases, *Ri*CE2 and *Ri*CEX, from the Firmicute *Roseburia intestinalis,* which together deacetylate complex galactoglucomannan (GGM). The 3D-structure of *Ri*CEX with a mannopentaose in the active site shows that the CBM35 domain of *Ri*CEX forms a confined complex, where the axially oriented C2-hydroxyl of a mannose residue points towards the Ser41 of the catalytic triad. Cavities on the *Ri*CEX surface may accept galactosylations at the C6 positions of mannose adjacent to the mannose residue being deacetylated (subsite −1 and +1). In depth characterization of the two enzymes using time-resolved NMR, HPLC and mass spectrometry demonstrates that they work in a complementary manner. *Ri*CEX exclusively removes the axially oriented 2-*O*-acetylations on any mannose residue in an oligosaccharide, including double acetylated mannoses, while the *Ri*CE2 is active on 3-*O-,* 4-*O-* and 6-*O-*acetylations. Activity of *Ri*CE2 is dependent on *Ri*CEX removing 2-*O*-acetylations from double acetylated mannose. Furthermore, transacetylation of oligosaccharides with the 2-*O* specific *Ri*CEX provided new insight to how temperature and pH affects acetyl migration on mannooligosaccharides.

**Significance statement:** Acetylations are an important feature of hemicellulose, altering the physical properties of the plant cell wall, and limiting enzyme accessibility. Removal of acetyl groups from beta-mannan is a key step towards efficient utilization of mannans as a carbon source for gut microbiota and in biorefineries. We present detailed insight into mannan deacetylation by two highly substrate-specific acetyl-mannan esterases (AcMEs) from a prevalent gut commensal Firmicute, which cooperatively deacetylate complex galactoglucomannan. The 3D structure of *Ri*CEX with mannopentaose in the active site has a unique two-domain architecture including a CBM35 and an SGNH superfamily hydrolytic domain. Discovery of mannan specific esterases improves the understanding of an important step in dietary fiber utilization by gut commensal Firmicutes.

## Introduction

Enzymatic acetylation and deacetylation are involved in numerous processes in Nature. N-acetylation of lysine in histones is crucial for the control of gene transcription (1); acetylation or deacetylation of N-acetylglucosamine and N-acetylmuramic acid in microbial peptidoglycan affects its degradation by lysozyme (2). The efficacy of penicillins is also linked to acetylations, e.g. penicillin resistance caused by bacteria expressing cephalosporin esterases (3). Acetylations are important constituents in some of the most abundant polymers on earth like chitin, pectin, xylan and β-mannan (4, 5). Enzymes that process these polymers have adapted to a wide array of structurally diverse targets. Studying the structure-function relationships of acetyl esterases and transferases involved is shedding light on this important and very common covalent modification.

β**-**Mannans are polysaccharides present in the secondary cell walls of plants with a backbone consisting of *β*-1,4-linked D-Man*p* units (Fig. 1). The backbone can be interspersed with *β*-1,4 linked D-Glc*p*, as in konjac glucomannan (*Amorphophallus konjac*), decorated with *α*-1,6-D-Gal*p* as in carob galactomannan (*Ceratonia siliqua*), or contain both glucose and galactose such as the GGM in *Aloe vera* and Norway spruce (*Picea abies*) (6-8). Furthermore, mannans may contain substantial amounts of 2-*O*-, 3-*O-*, and 6-*O*-acetylations on any mannose residue in the chain, as well as 4-*O*-acetylations (rare) on the non-reducing end of the chain.

**Fig. 1.**
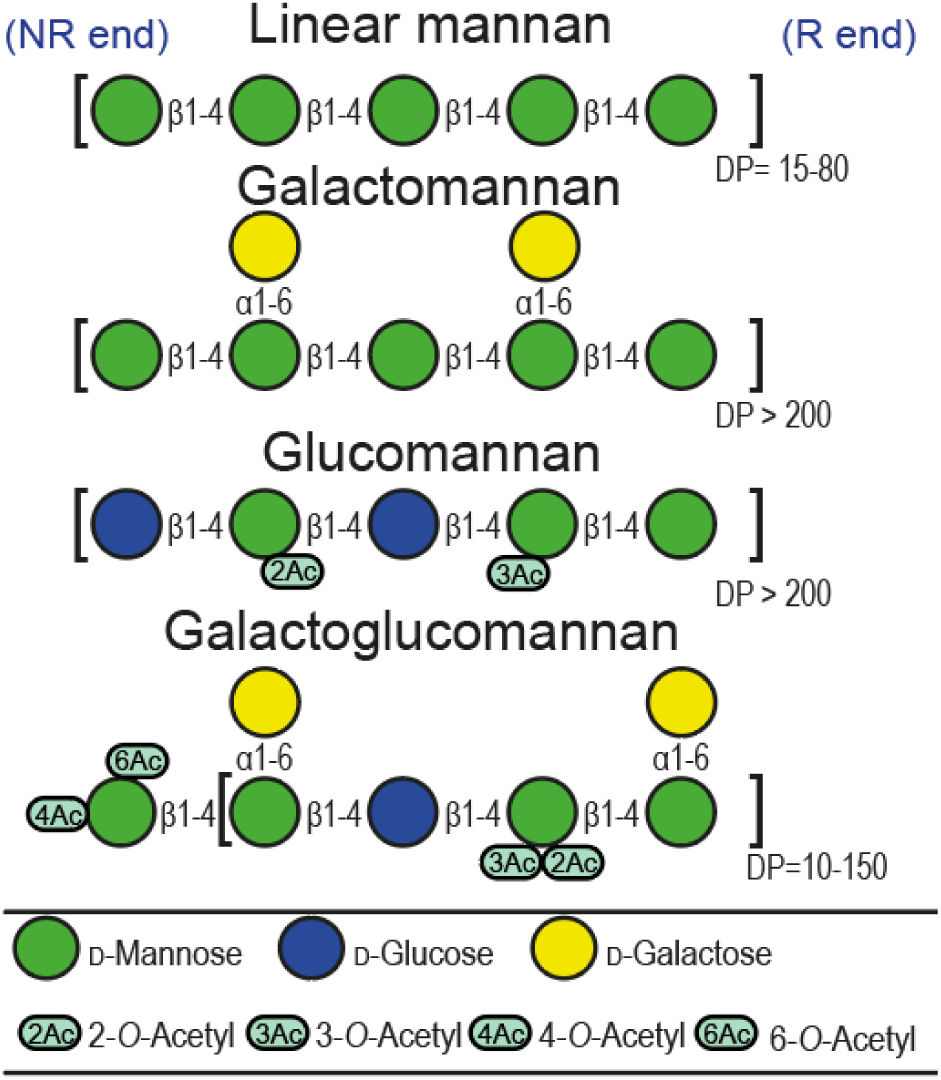
A cartoon presentation of various types of mannan found in plants (figure is an adaptation from Fig. 1 in 20). Glucomannans like konjac glucomannan generally contain a low degree of acetylation (DA =0.07). Galactoglucomannans contains various DA’s, the spruce mannan used in this study (DA = 0.35) and Aloe vera mannan (DA > 0.5).

Depending on the plant origin, mannans differ in their monosaccharide composition, degree and location of acetylations (9). A unique characteristic of the 2-*O*-acetyl groups of mannans is their relative orientation: 2-*O*-acetylations are axial, as opposed to other common structural polysaccharides, like xylan, pectin and chitin, that all have acetylations in the equatorial plane of the sugar ring.

Acetylation of hemicelluloses affects their biological functions and a range of physicochemical properties and is thought to be a defense mechanism developed as a part of an evolutionary arms race against pathogens (5) as well as contributing to aggregation of cellulose microfibrils (10). Acetylations affect the solubility of the polysaccharides by restricting the formation of hydrogen bonds between chains in solution. Effects of mannan acetylations on the viscosity of solutions make them effective thickeners and stabilizers (9, 11) in the food and feed industry where mannans such as guar gum and konjac are commonly used (12, 13). Furthermore, the immunostimulatory properties of *Aloe vera* (AV) mannan, a common ingredient in nutraceuticals and cosmetics, have been linked to its high degree of acetylation (8). In the context of biorefining, acetylations reduce fermentability (14) and enzymatic degradability, and thus pose a limitation on the utilization of mannans as a feedstock for fermentation. In this respect, deacetylation of mannan by chemical pretreatment or enzymatic treatment with acetyl esterases is known to improve fermentation yields. On the other hand, the recalcitrance resulting from the presence of acetylations also restricts the number of gut microbes able to consume hemicellulose. This may be beneficial as means of providing selective growth of bacteria when hemicellulose is considered as an ingredient of food and feed.

Gut microbes are highly adapted to fermenting diet-derived complex carbohydrates and have evolved complex machineries called polysaccharide utilization loci (PULs) to utilize these carbohydrates (15, 16). Bacteroidetes and Firmicutes are the main polysaccharide degrading commensal bacteria in the human gut (17, 18). β-mannans constitute a small but significant part of human diet that can be found in a wide range of different food products. This may explain why mannan degradation is identified as one of the core pathways in the human gut microbiome (19). Some species of Firmicutes, such as *Roseburia intestinalis*, have developed sophisticated enzymatic machineries for complete β-mannan degradation; these PULs quite frequently contain esterases (20).

The esterases deacetylating xylan and mannan characterized so far have been classified together as acetylxylan esterases (EC 3.1.1.72). Carbohydrate esterases (CEs) are common in bacterial and fungal genomes and are often found in PULs together with glycoside hydrolases (GH) (20, 21). This highlights their biological role as accessory enzymes facilitating the processing of glycans. CEs are classified into 16 families according to the CAZy database (22), in which members of families 1-7 and 16 have been reported to be active on hemicelluloses in plant biomass. All CEs, with the exception of family CE4, are serine-histidine hydrolases (5). Esterases from the CE2 family share a two-domain architecture consisting of a catalytic SGNH hydrolase superfamily domain with a conserved catalytic dyad, and an accessory jelly roll domain (23). The two-domain structure results in an open active site able to accommodate a variety of substrates. So far, functionally characterized CE2 esterases have been largely non-specific; activities has been reported on 2-*O*-, 3-*O*- and 4-*O*-acetylated xylan, konjac glucomannan (24), and 6-*O*-acetylated mannan (5). Some CE2 esterases have been considered specific to 6-*O-* acetylations, which is the position they transacetylate when using vinyl acetate as the acetate donor (25). Enzymatic deacetylation of mannans has previously been described in literature (26, 27), but no esterase exclusively active on mannan has been reported.

Here, we report a detailed functional characterization of *Ri*CE2 and *Ri*CEX - two acetyl mannan esterases (AcMEs) from *R. intestinalis*. The esterases have complementary substrate specificities and act in tandem to deacetylate glucomannan and galacto(gluco)mannan. Furthermore, we report the first structure of an AcME with a mannopentaose co-crystallized in the active site, revealing how the two-domain architecture of *Ri*CEX facilitates specificity towards the axially oriented 2-*O*-acetylation. To the best of our knowledge, this structure is also the first carbohydrate esterase of any kind crystallized with a substrate in the active site.

## Results and discussion

A gene cluster in *R. intestinalis* encoding mannan degrading enzymes (20) contains two acetyl esterases, *Ri*CEX and *Ri*CE2. *Ri*CE2 is a 349 amino acid CAZy family 2 esterase which shares 29.46% identity with the xylan esterase CjCE2C (also known as CE2C;CJA_2889 and Axe2B) of *Cellvibrio japonicus* (Uniprot accession no B3PC75) (24). Analysis of the *Ri*CE2 sequence with InterProScan (28) identified the C-terminal residues 163-375 in *Ri*CE2 as an SGNH hydrolase domain, while residues 1-159 constitute a galactose binding superfamily domain. This two-domain architecture is characteristic for CE2 family esterases (23). BlastP searches (29) of *Ri*CEX (372 amino acids) returned very low sequence homology to any characterized esterases (20% sequence similarity with acetyl xylan esterase Axe2 of *Geobacillus stearothermophilus* (30). Despite the low similarity, the sequence still appeared to be a promising esterase candidate, due to its location in the *R. intestinalis* mannan PUL, and the N-terminal SGNH hydrolytic domain. Contrary to the CE2 family esterases, where the SGNH domains are present in the C-terminal, the *Ri*CEX hydrolytic domain is present in the N-terminal residues 35-209. InterProScan (28) was not able to predict a function for the 163 amino acid C-terminal domain of *Ri*CEX. This previously uncharacterized sequence organization captured our interest for further structural exploration.

Crystal structures for the *Ri*CEX with acetate (PDB ID code 6HFZ, 1.75 Å resolution) and mannopentaose (PDB ID code 6HH9, 2.4 Å resolution) in the enzyme active site reveal several structural feauteres. For the structure with mannopentaose, four mannose units were observed in the electron density maps. The enzyme consists of two domains connected by a linker of about 20 amino acids (Fig. 2A). The N-terminal domain (ca. 210 amino acids) displays an α/β-hydrolase (SGNH-hydrolase) fold while the C-terminal domain (ca. 140 amino acids) comprises a jelly-roll beta-sandwich fold. Structural alignment of the complete *Ri*CEX structure against the Protein Data Bank using PDBeFold (31) returned hits with low scores, indicating this is a hitherto unidentified fold. The two structures with the highest scores were Ape1, a peptidoglycan *O*-acetylesterase from *Neisseria meningitidis* (Fig. 2B) (PDB ID code 4K40, Q-score 0.17) (32), and the *Cj*CE2B (Uniprot accession no. F7VJJ8) carbohydrate esterase from *C. japonicus* (Fig. 2C) (PDB ID code 2W9X, Q-score 0.12) (24). It is apparent that the second domain of these three enzymes adopts different positions in space when aligning their catalytic SGNH domains (Fig. 2A-C). These differences are not due to domain flexibility. For the Ape1 enzyme, the second domain is inserted in the middle of the SGNH domain sequence and is kept in place by two short linkers consisting of 3-4 amino acids. The second domain in the *Cj*CE2B enzyme, located at the N-terminal end of the SGNH domain, also adopts a very different position than that found for the C-terminal domain in *Ri*CEX, connected to the SGNH domain via an eight-residue linker.

**Fig. 2.**
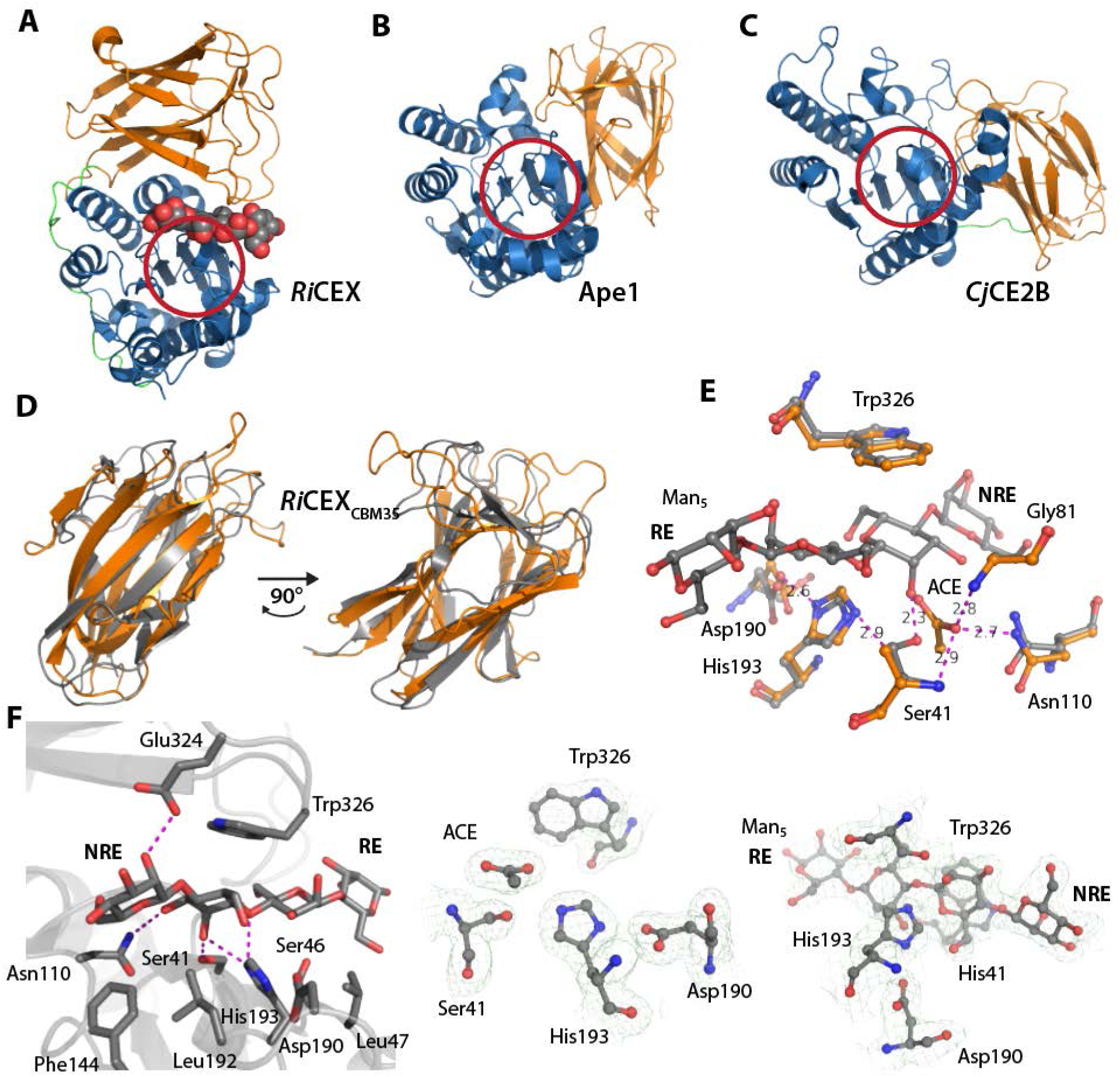
Comparison with *Ri*CEX homologues and structural details of *Ri*CEX. (A) Crystal structure of *Ri*CEX, with four of the mannose residues present in mannopentaose (spheres) visible in the electron density map. The catalytic domain (blue) is connected to the CBM35 domain (orange) by a linker region (green). Panels (B) and (C) display the structural homologs Ape1 and *Cj*CE2B that are closest to *Ri*CEX in the PDB database, respectively. The catalytic domains (blue) in (B) and (C) are oriented to superimpose with the *Ri*CEX catalytic domain in A). The red rings in panels (A-C) indicate where the residues of the catalytic triad are located. In panel (D) the *Ri*CEXCBM35 domain (orange) is superimposed on the *Ct*CBM35 domain (grey) and shown at two rotations, demonstrating the highly similar folds. (E) Display of the *Ri*CEX superposed with acetate (orange carbons, ACE denotes acetate) or mannopentaose (grey carbons) in the active sites. In both structures Asp190 form a hydrogen bond with His193 (2.6 Å). In the acetate bound structure the His193-N^ε2^ is hydrogen bonded with the Ser41-OH O-atom (2.9 Å). The O-atom of the acetate molecule pointing towards the oxyanion hole is hydrogen bonded by Ser41-N (2.9 Å), Asn110-N^γ 2^ (2.7 Å) and Gly81-N (2.8 Å), indicated by dotted magenta lines to the acetate molecule. In the mannopentaose bound structure, the hydrogen bond between His193-N^ε 2^ and the Ser41-OH O-atom is replaced with a Ser41-OH O-atom hydrogen bond to the C2-OH group of the mannose residue in the esterase catalytic site (2.3 Å), implying that the Ser41 side chain change conformation during catalysis. (F) Several charged or polar residues form hydrogen bonds to the mannopentaose ligand (Ser41, Asn110, His193 and Glu324). Ser46 may interact with the ligand through a water bridge. The hydrophobic residues Leu47, Phe144, Leu192 and Trp326 are part of the ligand-protein contact surface. Trp326 plays an important role, stacking on top of the mannose residue in the active site. The electron density maps of the active sites are shown to the right in panel (F). The non-reducing end (denoted NRE) and the reducing end (denoted RE) are labelled in the mannooligosaccharide ligand to aid interpretation of the figures.

Structural alignment searches using the C-terminal, non-catalytic domain of *Ri*CEX, identified a carbohydrate binding module 35 (CBM35), similar to that of *Clostridium thermocellum* (Fig. 2D) (PDBid 2W1W, Q-score 0.39)(33). CBM35 modules have previously been demonstrated to bind decorated mannans (34). The *Ri*CEXCBM35 domain appears to be involved in substrate recognition and binding, forming a lid over the SGNH-domain active site (Fig. 2). Several hydrophobic amino acids, Ile79, Phe86, Ala89, Ile263, Leu330, Leu332 and Ile367, contribute to stabilize the inter-domain interaction together with the three pairs of charged or polar amino acid residues Glu119-His338, His116-Glu337 and Gln85-Thr334 (Fig. 2).

In the active site of the SGNH domain, three residues Ser41, His193 and Asp190 form the hydrogen-bonded catalytic triad, and the amide nitrogen atoms of Ser41 and Gly81 and the N^γ2^ of Asn110 line the oxyanion hole (Fig. 2E). The high 2-*O*-acetylation specificity of *Ri*CEX can be explained by the ligand bound structure. Amino acids from both the SGNH and CBM35 domains form specific interactions with the mannopentaose and aligns it such that the C2-OH group is only 2.3 Å from the Ser41-OH group (O-O distance), close to the oxyanion hole, which forms a cavity that fits an acetyl group. In the apo-*Ri*CEX structure we can clearly see an acetate molecule in the oxyanion hole (Fig. 2E) and this arrangement could represent a product complex. Such binding of acetate in absence of carbohydrate substrate has also been observed for Ape1 (32). The binding of acetate indicates how intermediates in the reaction may be positioned and provides structural insight useful for further mechanistic studies.

Fig. 2F shows interactions between *Ri*CEX and the mannopentaose ligand. Two amino acids from the *Ri*CEX_CBM35_ domain contribute to ligand binding, namely Glu324 and Trp326. The Glu324 forms a hydrogen bond to the *O*2 hydroxyl group of the mannose at the non-reducing end next to the active site (subsite −1 according to Hekmat et al (35)). Aromatic amino acids are often involved in protein-carbohydrate interactions, and in *Ri*CEX the Trp326 amino acid side chain stacks on top of the mannose unit in the active site. The CBM35 appears to play a central role in reactivity of the enzyme. A truncated version of the enzyme, containing only the catalytic domain, was not active either on natural substrates, or pNP (*p*-Nitrophenol or 4-Nitrophenol) acetate Fig. S1). An observed lack of activity of the SGNH domain indicates that it does not have an inherent esterase activity alone, but rather has adapted to 2-*O*-deacetylation by interaction with the CBM35 domain. The activity appears to depend on the two domains forming a confining clamp that provides substrate interactions and aid protein stability. This hypothesis was further supported by the results of thermal shift assay (Fig. S2), which showed a decrease in stability for the SGNH domain (as compared with the complete *Ri*CEX), and a lack of folding for the CBM35 domain. Interestingly, the two-domain arrangement results in cavities or clefts on each side of the active site indicated by the two green and magenta patches in Fig. 3A. Both these cavities are arranged so that galactose residues linked to the mannose backbone (corresponding to −1 and +1 subsites) at the *O-*6 position would point towards these spaces. This feature may be important for recognizing galactose decorated mannans and may also influence the substrate specificity of the enzyme.

**Fig. 3.**
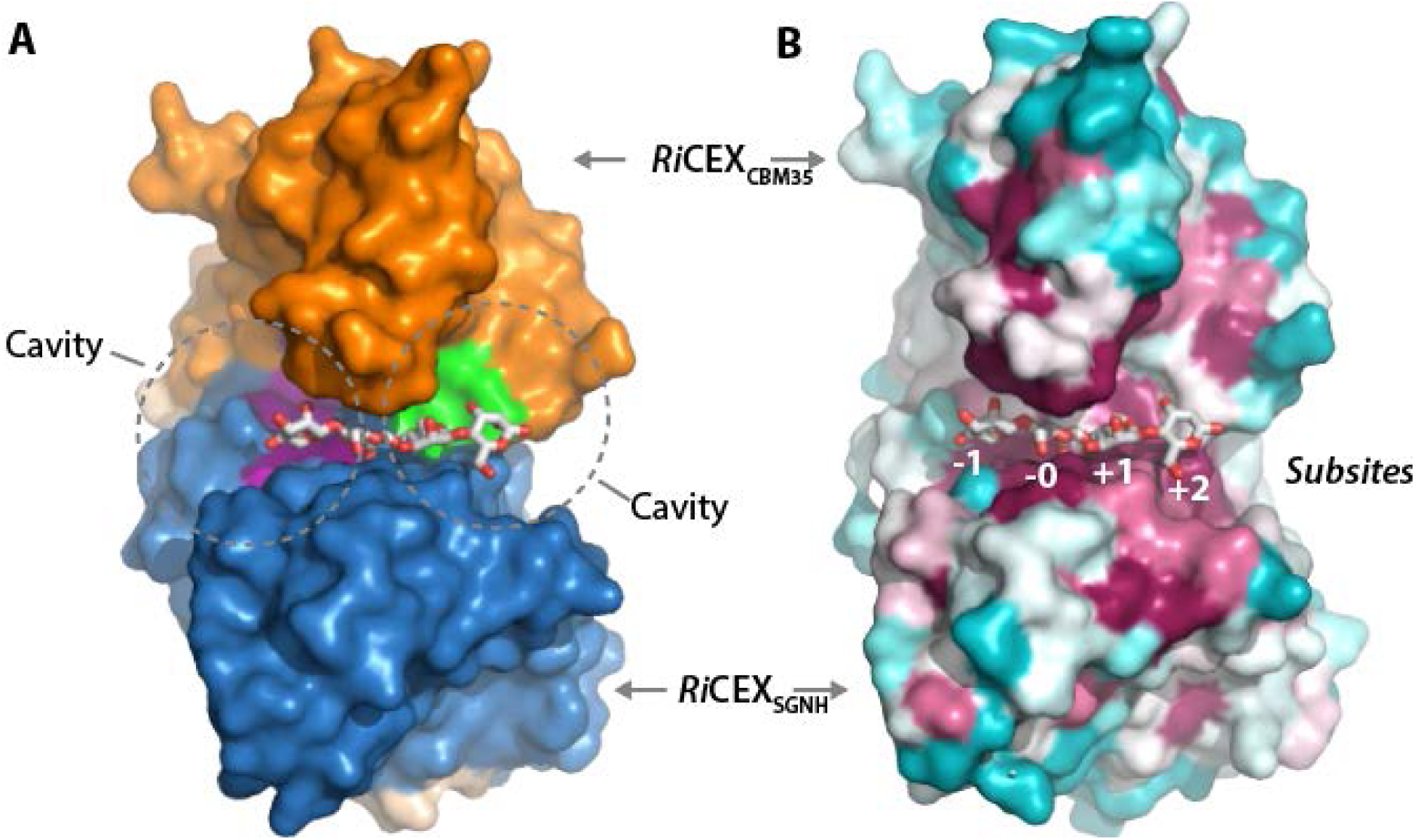
The 3D structures of substrate and product complexes of *Ri*CEX. (A) the magenta and green patches in the interface between the *Ri*CEX catalytic domain and the *Ri*CBM35 domain indicate cavities on the enzyme surface that may bind galactose residues decorating galactoglucomannan at the C6 position. In panel (B) the conservation score derived by ConSurf is projected on the *Ri*CEX surface. The magenta patches indicate highly conserved residues, which are concentrated around the substrate binding site. Subsites are labelled below the substrate according to the nomenclature suggested by *Hekmat et al (35).*

Since mannan degradation is a common process in many microbial ecosystems, we searched for microbes carrying protein sequences related to *Ri*CEX. Searching the UniProtKB database with a Hidden Markov Model (HMM, detailed description in the supplementary information) of both domains of *Ri*CEX, we identified 449 protein sequences with the same two-domain architecture. Sequence alignment using MView showed that the catalytic triad (Ser41, His193 and Asp190) and the Trp326 of the CBM domain were conserved in the 90% consensus sequence (Fig. S3). While a vast majority of *Ri*CEX homologues came from the phylum *Firmicutes* (435), 16 hits were identified in *Bacteroidetes.* Analysis of the neighbouring genes in both Firmicutes and Bacteroidetes genomes and alignment of *Ri*CEX along with five of its homologues (Fig. S4) from taxonomically distant bacterial taxa suggests that the active site arrangement might be adapted to a wider range of roles than just deacetylation of β-mannans.

To assess if the conserved residues correspond to functionally important structural features of *Ri*CEX, a subset of 77 sequences was selected from the HMM results by using a more stringent E-value, and the conservation pattern was projected on the *Ri*CEX protein surface using the ConSurf Server (Fig. 3B) (36). Highly conserved amino acid residues (magenta), semi-conserved (white) and variable residues (blue) are indicated, and it is apparent that both the SGNH and CBM35 domains contribute with conserved residues around the substrate binding site. The residues Ser41, Asn110, His193, Leu47, Phe144, Leu192 of the of SGNH domain and Trp326 of the CBM35 domain are all conserved and are interacting with the substrate in the active site.

The mannan PUL in *R. intestinalis* encodes an extracellular mannanase -*Ri*GH26 which breaks down mannan into oligosaccharides, which are then internalized via an ABC transporter deacetylated by the esterases and further metabolized (20). To approximate these conditions, mannans used in activity testing were hydrolyzed with *Ri*GH26 prior to deacetylation. The two esterases were active on a wide range of mannans but not on structurally similar substrates such as acetylated xylan, cellulose monoacetate or chitin oligosaccharides (Fig. S5). A selection of mannose-based substrates was used to examine the impact of galactosylations (Norway spruce mannan), mannose units with multiple acetylations (*Aloe vera* mannan) and different acetylation distributions (konjac mannan, and its chemically acetylated version) on enzyme activity. Neither *Ri*CE2 nor *Ri*CEX were active on birch xylan (as well as its hydrolysate produced with a GH10 xylanase from *R. intestinalis* (21)), cellulose monoacetate or chitopentaose (Fig. S5D and E). The deacetylation specificity was the same on all mannans: there was a partial deacetylation seen whenever one of the esterases was used on its own (red and blue traces Fig. 4A and S5A-C and F). When the two esterases were used together, a complete deacetylation was observed (purple traces Fig. 4A and S5A-C and F).

**Fig. 4.**
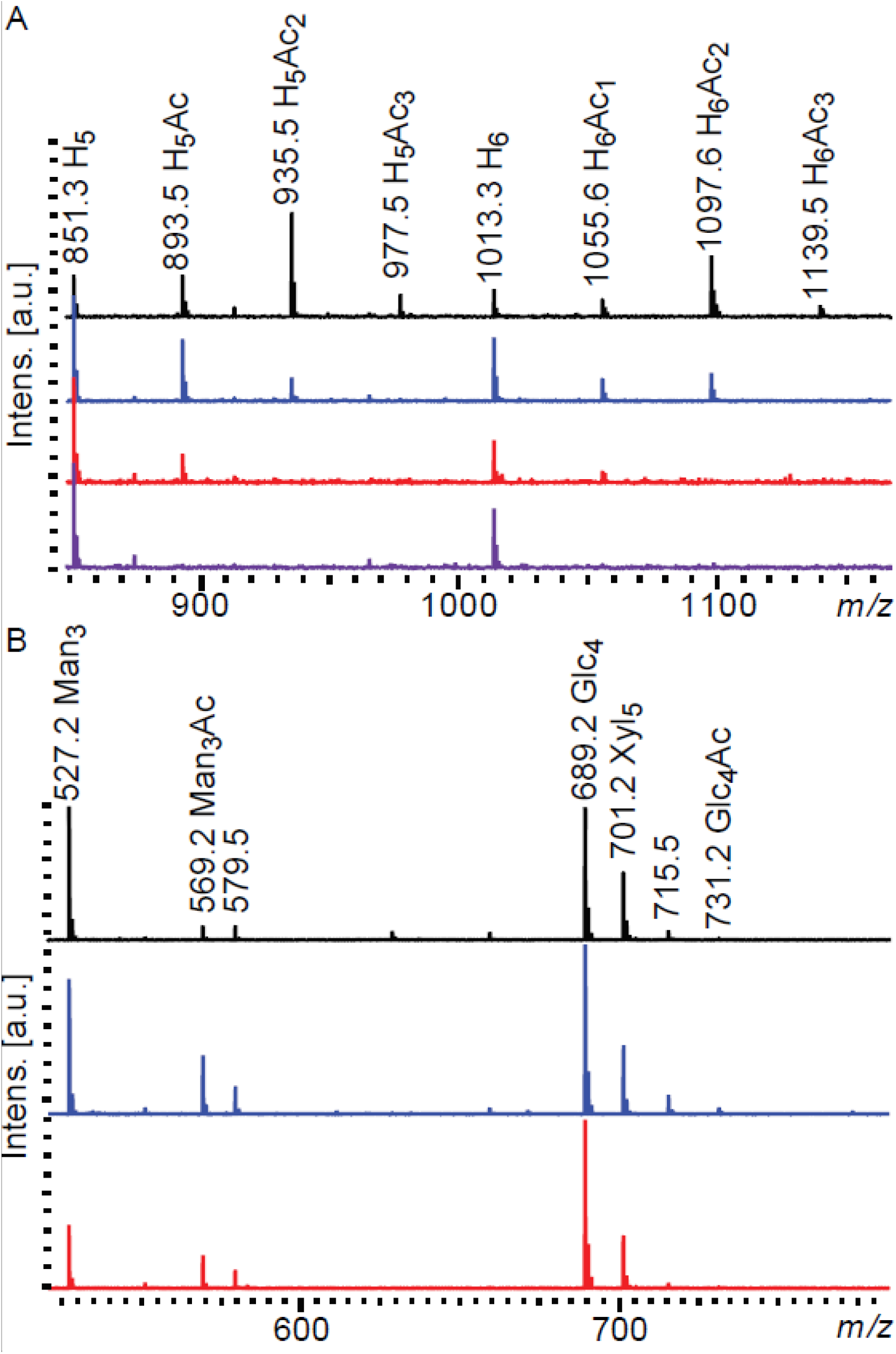
MALDI-ToF spectra of deacetylation and transesterification reactions. (A) 10 mg/mL Spruce GGM hydrolysate (untreated in black) was treated with *Ri*CE2 (in blue), *Ri*CEX (in red) and the two enzymes in combination (purple). When either of the esterases was used, a decrease in the intensitites of peaks signifying acetylated oligosaccharides was observed. Using the two enzymes in combination lead to a complete deacetylation of spruce mannan. (B) Transesterification of a mixture of 1 mg/mL of mannotriose, cellotetrose and xylopentose with a mixture of vinyl acetate, propionate and butyrate. No spontaneous esterification was observed (black trace). Out of all possible substrate combinations, both *Ri*CE2 (in blue) and *Ri*CEX (in red) produced acetylated mannotriose as the primary product. *Ri*CE2 produced a minor amount of acetylated cellotetraose (*m/z* 731) as a by-product. H-hexose, Glc-glucose, Man-mannose, Xyl-xylose, Ac – acetylation.

Commercially available and in-house produced substrates contain a wide range of acetylations present in a variety of positions on a mannose unit, and different locations in the oligosaccharide chain, making it challenging to precisely determine the specificity of esterases. In transesterification reactions, non-acetylated substrates are treated with esterases in the presence of an appropriate acyl donor, resulting in an amount of highly specific acetylated oligosaccharides consistent with the free energy of the reaction. Transesterification with vinyl acetate has been used as a measure of substrate specificity of acetyl esterases before (37). To further explore the substrate specificity of *Roseburia* esterases, we used vinyl-acetate, -propionate and -butyrate as acyl donors. Both enzymes were able to transacetylate and transpropylate (although to a much lesser extent than acetylate) but not transbutyrylate mannotriose (Fig. 4 and S6). Both esterases were able to transacetylate all mannose-based substrates tested, including mannooligosaccharides with DP range 2-6, 6^1^-α-D-galactosyl-mannotriose, 6^3^, 6^4^-α-D-galactosyl-mannopentaose, as well as hydrolysates of AV mannan and Norway spruce GGM produced in house (Data not shown).

As a means of demonstrating specificity transacetylation experiments were performed on a mixture of mannotriose, cellotetraose and xylopentaose, with vinyl-acetate, -propionate and -butyrate as ester donors. Acetylated mannotriose was the only product observed in the MALDI ToF spectra (Fig. 4B), clearly indicating the mannan acetylesterase specificity of *Ri*CEX and *Ri*CE2.

The partial deacetylation observed in Fig. 4 and S5 warranted more detailed analysis of the deacetylation preferences of the esterases. NMR analysis (Fig. 5) revealed that the partial deacetylation seen in *Ri*CEX-treated samples was due to the fact that *Ri*CEX exclusively removed all 2-*O*-acetylations from AV and Norway spruce GGM (Fig. 5B and 5F, respectively). *Ri*CE2 removed the 3-*O*- and 4-*O*-acetylations as well as some of the 6-*O*-acetylations from single acetylated mannose units in the same substrates (Fig. 5C and 5G). Importantly, in the AV samples which contained a high number of double acetylated mannose, *Ri*CE2 was not able to remove the 3-*O*- and 6-*O*-acetylations present on the same mannose as a 2-*O*-acetylation (Fig. 5C). Adding *Ri*CEX to the solution of AV after the *Ri*CE2 enzymatic treatment and vice versa lead to a complete deacetylation of both substrates (Fig. 5D and 5H), indicating that the two esterases have interdependent specificities, and are both required for complete deacetylation of mannans.

**Fig. 5.**
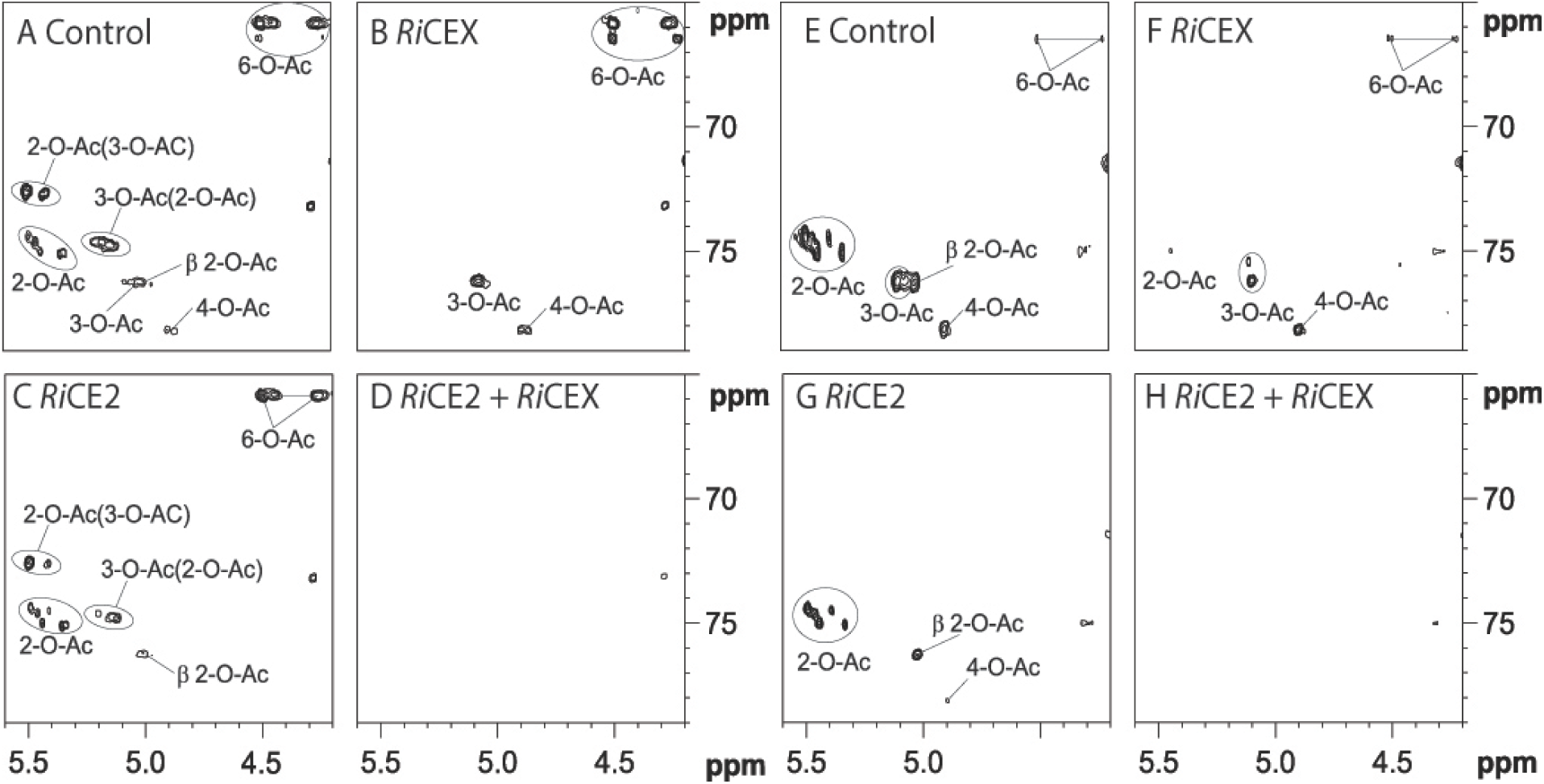
2D ^13^C HSQC NMR spectra for area of interest of AV mannan *Ri*GH26 hydrolysate (panels A-D) and Norway spruce mannan *Ri*GH26 hydrolysate (panels E-H) treated with the esterases. (A) AV hydrolysate without enzyme addition, contained a high degree of acetylation on all possible positions, and a high degree of double acetylation. (B) When treated with *Ri*CEX, the peaks corresponding to 2-*O*-acetylations disappear, and the shift values for peaks corresponding to 3-*O*- and 6-*O*-acetylations change due to the removal of 2-*O*-acetylations from double acetylated mannoses. (C) Treatment with *Ri*CE2 removed all the non-reducing end 4-*O*-acetylations, the majority of 3-*O*- acetylations, and some of the 6-*O*-acetylations. (D) Treatment with both esterases at the same time removed all acetylations. (E) Spruce mannan hydrolysate without enzyme addition contained prevalently 2-*O*-, some 3-*O*-acetylations, and a lower degree of 4-*O*- and 6-*O*-acetylation. (F, G) Both enzymes exhibited similar activity on spruce mannan as on AV. (H). Acetylations in parenthesis signify the adjacent acetylations on double acetylated mannose (2-*O*-Ac(3-*O*-Ac) - -2*-O*-acetylation adjacent to a −3-*O*-acetylation).

*Ri*CE2 is a broad specificity esterase capable of removing 3-*O*-, 4-*O*- and 6-*O*-acetylations, although its activity is limited by the presence of 2-*O*-acetylations as one of multiple acetylations on a single mannose. This dependence of *Ri*CE2 on *Ri*CEX was also observed in the reaction rates calculated from time-resolved NMR analysis of reactions with AV as the substrate. Reaction rates showed that *Ri*CE2 was much slower when 2-*O*-acetylations were present (Table 1). The presence of 3-*O*- and 6-*O*-acetylations on the same mannose residues containing 2-*O*-acetylation did not affect the apparent k_cat_ of *Ri*CEX, while the turnover rate of *Ri*CE2 more than doubled when 2-*O*-acetylations were removed (Table 1). *In vivo*, the enzyme pair would act in complement, with the specialist *Ri*CEX removing the 2-*O*-acetylations that limit the activity of the generalist *Ri*CE2.

**Table 1.**
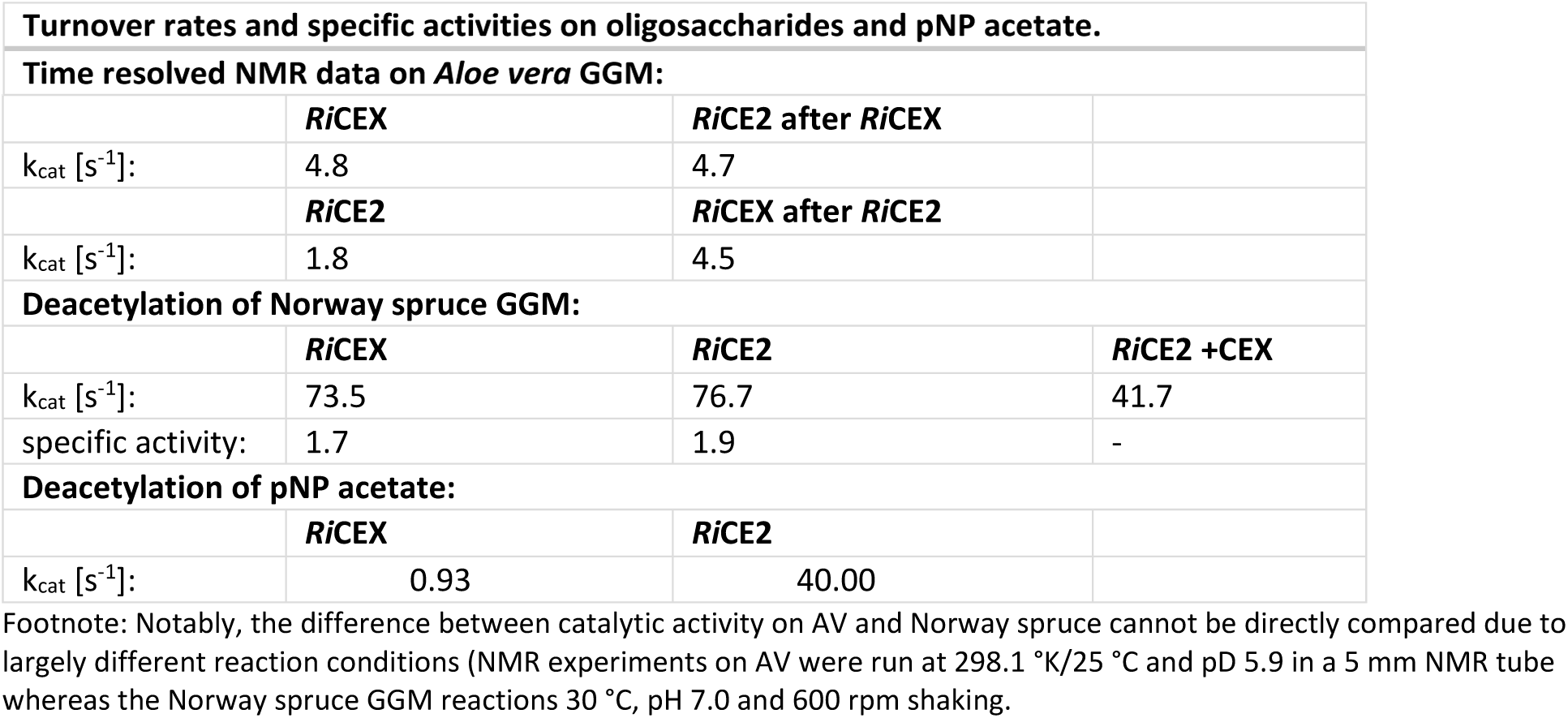
Reaction rates of *Ri*CEX and *Ri*CE2. Turnover rate in time-resolved NMR analysis of deacetylation determined based on the acetate released in the initial 15 minutes of reaction. Two identical samples of AV mannan hydrolysed with RiGH26 mannanase were prepared and treated with 1) 62.5 nM loading of *Ri*CE2 for 16 hours, at 298.1 °K, then 10 nM loading of *Ri*CEX 2) 10 nM loading of *Ri*CEX for 16 hours, then 62.5 nM loading of *Ri*CE2. Turnover rate and specific activity [nanomole acetate/s/µg enzyme] of esterases on Norway spruce GGM RiGH26 hydrolysate were determined based on the amounts of acetate released from samples of 100mg/mL of substrate in 30 minutes. Turnover rates of both esterases on pNP acetate appear lower than on oligosaccharides.

| Turnover rates and specific activities on oligosaccharides and pNP acetate. |
| --- |
| Time resolved NMR data on *Aloe vera* GGM: |

|  |  |  |  |
| --- | --- | --- | --- |
|  | ***RiCEX*** | ***RiCE2* after *RiCEX*** |  |
| $k_{cat}$ [$s^{-1}$]: | 4.8 | 4.7 | |
|  | ***RiCE2*** | ***RiCEX* after *RiCE2*** |  |
| $k_{cat}$ [$s^{-1}$]: | 1.8 | 4.5 | |
| **Deacetylation of Norway spruce GGM:** |  |  |  |
|  | ***RiCEX*** | ***RiCE2*** | ***RiCE2* +CEX** |
| $k_{cat}$ [$s^{-1}$]: | 73.5 | 76.7 | 41.7 |
| specific activity: | 1.7 | 1.9 | - |
| **Deacetylation of pNP acetate:** |  |  |  |
|  | ***RiCEX*** | ***RiCE2*** |  |
| $k_{cat}$ [$s^{-1}$]: | 0.93 | 40.00 | |

Footnote: Notably, the difference between catalytic activity on AV and Norway spruce cannot be directly compared due to largely different reaction conditions (NMR experiments on AV were run at 298.1 °K/25 °C and pD 5.9 in a 5 mm NMR tube whereas the Norway spruce GGM reactions 30 °C, pH 7.0 and 600 rpm shaking.

Activity assays using para-Nitrophenyl (pNP) acetate, which is a commonly used substrate when screening for esterase activities showed that both esterases appear to be most active near neutral pH (Fig. 6). Notably, pNP acetate was a highly inadequate substrate for *Ri*CEX with the apparent turnover rate nearly 2 orders of magnitude lower than on Norway spruce GGM (Table 1). This is not surprising, considering the topology of the *Ri*CEX active site and the structural difference between the planar phenol ring of pNP and the axially oriented 2-O-acetylation on D-Man*p*; however, this result highlights the necessity for careful selection of substrates and methods when measuring esterase activity.

**Fig. 6.**
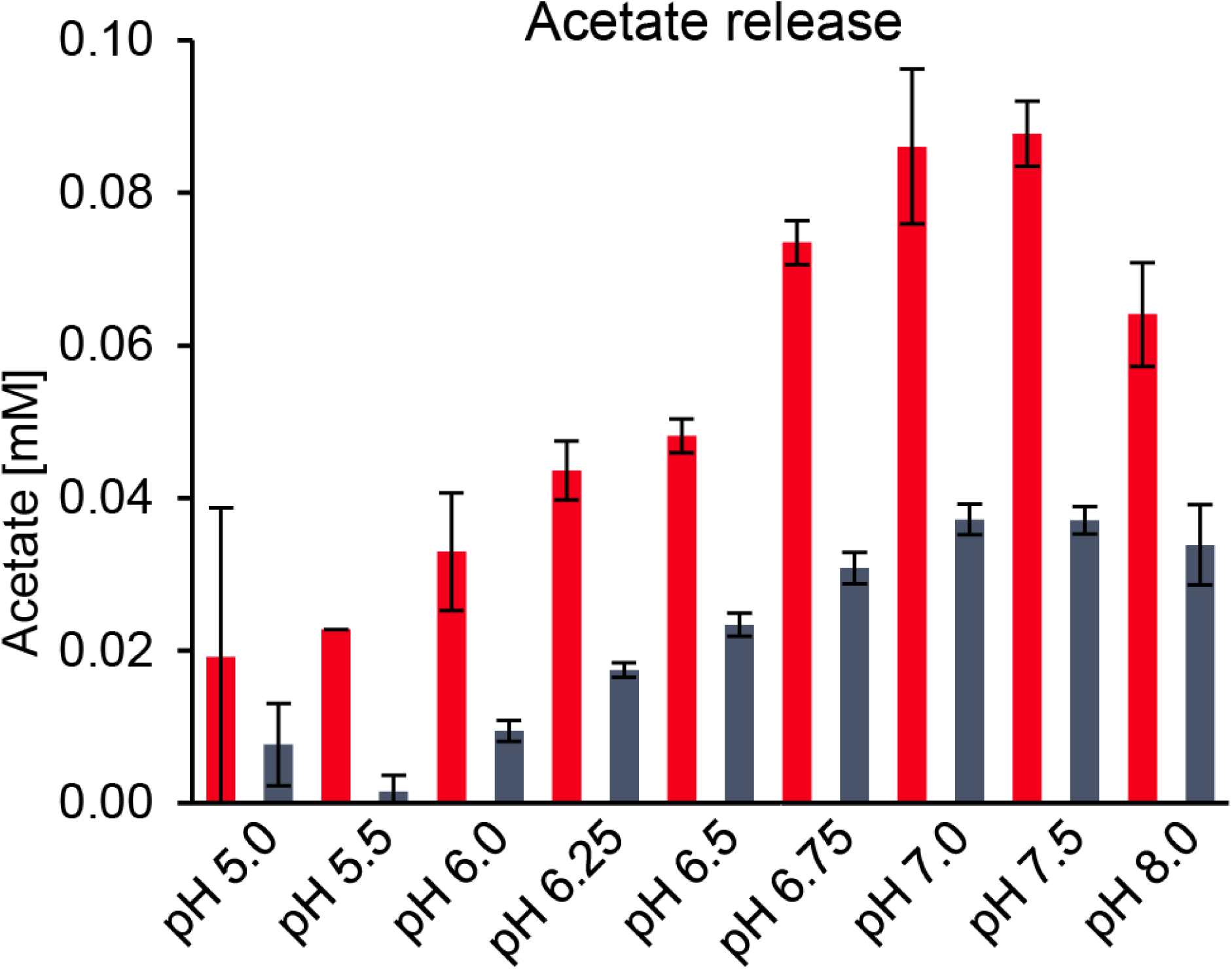
pH optima of RiCEX (in red) and RiCE2 (in blue) as determined by the amounts of acetate released form pNP acetate at different pH. Error bars indicate standard deviation between triplicates.

Both esterases showed near identical kcat and specific activities (Table 1) on *Ri*GH26 hydrolyzed Norway spruce GGM substrate. In deacetylation rate measurements at pH 7.0 and 35 °C (approximating intracellular conditions of a gut commensal bacterium), at equal loadings (25nM final concentration) of each esterase the apparent total rate of deacetylation was lower than when 50nM of either esterase was used on its own. The lower resulting rate of deacetylation suggests that the esterases were competing for substrate, an effect observed previously in lignocellulose degrading enzyme cocktails (38). Reaction rates obtained in time-resolved NMR showed an increase in the *Ri*CE2 activity after removal of 2-*O*-acetylations. This effect was not observed in the activity on Norway spruce GGM (Table 1), since the substrate used had much fewer double acetylated mannose units.

There is a growing body of evidence that acetylations on hemicellulose are prone to migration on sugar rings and also across glycosidic linkages (39, 40). This phenomenon requires particular attention when characterizing acetylesterases, since the distribution of acetylations will change under certain conditions. Elevated pH and temperature (>90°C) are known to induce acetyl migration on mannose and xylose (39, 41, 42). In experiments conducted on monosaccharides, it has been shown that the acetyl groups migrate at pH > 6.0. In both D-glucose and D-galactose, acetylations initially present in the 2-*O*-position appear to move in a ‘clockwise’ (2-*O*-→3-*O*-→4-*O*) direction at pD >7 (43, 44), while at pD <3.0 D-galactose was shown to deacetylate without the acetyl group migrating (44). Notably, we observed that migration happens also on the polymers and at even less severe conditions. This phenomenon changes the properties of mannans as substrates and as biorefining feedstocks. At pH 5.9 and 20°C *Ri*CEX transacetylated all three Man*p* units of mannotriose, exclusively in the 2-*O*-position (Fig. S7 A) without any observed migration occurring. Exposure to just 60°C for one hour at pD 5.9 (Fig. S7 B), incubation at pD 7.4 at room temperature (Fig. S7 C), or incubation pD 7.4 at 60°C (Fig. S7 D), caused a decrease in the signals for 2-*O* acetylations and an appearance of signals for 3-*O*-, 4-*O*-and 6-*O*-acetylations, the latter two only in the non-reducing end mannose. At these conditions, the acetylations on the non-reducing end migrated in the same ‘clockwise’ 2-*O*- → 6-*O*-direction as described before on Gal*p* monosaccharides (41). The glycosidic bond prevents migration from 3-*O*-→ 6-*O*- and thus limit the migration from 2-*O*-/3-O-→6-*O*- for the reducing end and the intra-chain Man*p*. Migration pictured in Fig. S7 has progressed until an apparent equilibrium of acetyl distribution was reached, with a proportion of the acetylations remaining on the 2-*O-* position. Using the 2-*O*-deacetylation specificity of *Ri*CEX, we were able to produce a sample of Norway spruce GGM with all 2-*O-* acetylations removed. We examined the acetylation levels in solutions of Norway spruce GGM exposed to migration invoking conditions, then treated with *Ri*CEX to remove newly accumulated 2-*O-* acetylations. Both pH increase (Fig. S8) and heating to 60 °C for one hour (Fig. S9) induced a re-distribution of acetylations resulting in new 2-*O*-acetylations. Treatment with *Ri*CEX and repeated migration resulted in a complete removal of acetylations by only removing the 2-*O-* bound acetyls. This indicates that the acetyl migration does not proceed in a particular direction, but rather is a re-distribution process. A detailed description of acetyl migration is provided in the supplementary information.

## Concluding remarks

This study provides detailed insight into enzymatic deacetylation of β-mannans by the abundant human gut commensal bacterium *R. intestinalis*. The deacetylation apparatus of *R. intestinalis* consists of the highly specialized 2-*O*-acetylation specific *Ri*CEX, and *Ri*CE2 that removes all remaining types of acetylations. Both enzymes are necessary for deacetylation, and their activities are complementary. Both esterases have a two-domain structure with an SGNH superfamily hydrolytic domain and an accessory domain – a galactose binding superfamily domain in *Ri*CE2 and a CBM35 in *Ri*CEX.

Selective activity of *Ri*CEX was instrumental in our study of acetyl migration on mannooligosaccharides. We have demonstrated that acetyl migration proceeds towards the 2-*O*-position when the acetylations in that position are removed from an oligosaccharide. Our results show explicitly that the choice of substrate and reaction conditions is crucial for studying the specificity of hemicellulose esterases.

The potential for industrial application of this enzyme pair is quite apparent. With the activity on gluco- and galacto(gluco)mannans, the pair could be applied to various mannan-containing feedstocks to reduce the recalcitrance of mannans, modify the viscosity of mannans in solution, or to rationally design oligosaccharides with distinct esterification patterns via specific deacetylation or transesterification.

## Acknowledgments

Cellulose monoacetate with a degree of acetylation of 0.6 was a kind gift from Qi Zhou (KTH, Stockholm). A chemically acetylated konjac glucomannan prepared according to Bååth et al. (45) was kindly provided by Francisco Vilaplana. Norwegian Research council grant no. 244259, 208674/F50, 226244, and 226247 supported this work.

## Materials and Methods

### Enzymes

Recombinant *Ri*CEX (ROSINTL182_05471) and *Ri*CE2 (ROSINTL182_05473) were produced in *Escherichia coli* BL21 Star (DE3) as described previously (46). Truncated versions of *Ri*CEX and seleno-L-methionine substituted *Ri*CEX were produced as described in SI Materials and Methods.

### Substrates

Galactoglucomannan from Norway spruce (*Picea abies*) and glucuronoxylan from Birch (*Betula pubescens*) were produced in house from dried wood chips. The wood was milled into <2mm particles, and steam exploded at 200°C (10 minutes reactor residence time) in 5-6 kg batches. The liquid soluble fraction containing the hemicelluloses was extracted by washing the steam-exploded material with MilliQ water in a 50 µm pore WE50P2VWR bag filter (Allied filter systems, England). The liquid fraction of hemicellulose was then filtered through a 5 kDa spiral wound Polysulphone/polyethersulphone polyester ultrafiltration membrane (GR99PE, Alfa Laval, Denmark) using a GEA pilot scale filtration system Model L (GEA, Denmark). The fraction retained by the membrane was concentrated, collected and freeze-dried to become the Norway spruce GGM sample. Birch xylan was produced according to the protocol described in Biely *et al*. (47). 6^1^-α-D-Galactosyl-mannotriose; 6^3^, 6^4^-α-D-Galactosyl-mannopentaose, penta-N-acetylchitopentaose, mannotriose, mannotetrose, mannopentaose, konjac and carob mannans were purchased from Megazyme, Ireland. *Aloe vera* mannan (Acemannan) was purchased from Elicityl, France. Norway spruce GGM and *Aloe vera* mannan were hydrolysed with *Ri*GH26 in an unbuffered solution. Samples were then filtered through a pre-washed 1 mL Amicon Ultracel 3kDa ultrafiltration device (Merck KGaA, Germany) to remove the hydrolase and freeze dried.

### Activity analysis

Activities of the enzymes were tested by adding 1 µM of enzyme to 10 mg/mL solutions of carbohydrate substrates listed above. All substrate solutions were prepared with 20 mM sodium phosphate pH 5.9 to prevent acetyl migration. Reactions were incubated overnight in 20 mM sodium phosphate pH 5.9 at 30°C with 700 rpm shaking. In some analyses, MilliQ water was used instead of buffer to reduce background signals in MALDI-ToF MS as both enzymes were found to be active without buffers.

### Temperature and pH optima using 4-nitrophenyl acetate

In order to determine the pH and temperature optima for the enzymes, reactions with 0.5 mM pNP acetate were prepared using 50 mM sodium phosphate buffer (Sigma-Aldrich, Germany) in the pH range 5.5-8.0 (Sigma-Aldrich, Germany). pH optimum reactions were incubated at 25 °C. Due to the difference in deacetylation rate of pNP acetate by the two enzymes, 1 nM final concentration of *Ri*CE2 and 0.1 µM of *Ri*CEX were used in the pNP acetate experiments. Standard plots of 4-Nitrophenol (*p*-Nitrophenol) were prepared at each pH. To determine the optimum pH, 99 µL of sample mixture containing 1 mM of 4-nitrophenyl acetate in each of the buffer were added to the wells of a 96 well plate. 1 µL of enzyme solution was then added to the sample mixture and the reaction followed by measuring the absorbance at 405 nm at one minute intervals. All experiments were performed in triplicate, with two blanks for each condition set.

### HPLC Measurement of acetate release

Acetate content was analyzed on an RSLC Ultimate 3000 (Dionex, USA) HPLC using a REZEX ROA-Organic Acid H+ 300×7.8mm ion exclusion column (Phenomenex, USA) at 65°C, 10 µL injection volume, with isocratic elution using 0.6 mL/min of 5mM H2SO4 as mobile phase and a UV detector set to 210 nm.

### Crystallography

*Ri*CEX and *Ri*CE2 crystallization conditions were screened using several commercial high throughput 96 condition sitting drop screens, using 20 mg/mL, 10 mg/mL and 5 mg/mL solutions of protein in a 1:1 ratio with ready-mixed mother liquors (200 nL of each). Screening plates were set up using a mosquito HTS liquid handling robot (ttp labtech, UK).

Crystals were observed in spot G8 of the INDEX HT screen (Hampton Research, USA), containing 0.2M ammonium acetate, HEPES pH 7.5 and 25% w/v PEG 3350. Hanging drop optimization grids were manually set up with 0.2 M ammonium acetate, HEPES pH 7.0, 7.5 and 8.0, 20%, 25% and 30% w/v PEG 3350. Crystallization liquor was mixed 2 µL:2 µL and 1 µL:1 µL in the hanging drops, with additional 2 µL or 1 µL of 5 mg/mL solution of mannotriose, mannotetrose and mannopentose (Megazyme, Ireland) for co-crystallization.

Crystals were transferred to a cryo-solution containing mother-liquor with 35 % glucose before flash freezing in liquid nitrogen. Diffraction data were collected at beamlines ID23-1 and ID-29 at the European synchrotron Research Facility in Grenoble, France. For additional information on X-ray data collection, see supplementary material.

The initial structure was solved by single-wavelength anomalous diffraction (SAD) using selenomethionine to obtain an anomalous signal. Data was processed by XDS (48) and scaled by AIMLESS (49). The Phenix software package (50) was used to phase (AUTOSOL) (51) and build (AUTOBUILD) (52) the first structure. Subsequent structures were solved by molecular replacement (Phaser) (53) and refined using REFMAC (54). Model manipulations were carried out using Coot (55) and molecular graphics were generated using Pymol2 (Schrödinger).

### MALDI-ToF analysis

MALDI-ToF analysis of hydrolysis products was conducted on an UltraFlextreme MALDI-ToF instrument (Bruker Daltonics GmbH, Germany) equipped with a nitrogen 337 nm laser beam. Samples were prepared by mixing 2 μL of a 9 mg/mL solution of 2,5-dihydroxybenzoic acid (Sigma-Aldrich, Germany) in 30% acetonitrile (VWR) to an MTP 384 ground steel target plate (Bruker Daltonics GmbH, Germany) and 1 µL of hydrolysate (0.1-1 mg/mL). Sample drops were then dried under a stream of warm air.

Preparative HPLC, transesterification and NMR experiments are described in SI Materials and Methods.

