## Supplementary information for "A pair of esterases from a commensal gut bacterium remove acetylations from all positions on complex β-mannans"

**This PDF file includes:**

Supplementary text
Figs. S1 to S9
References for SI
Supplementary materials and methods

### Supplementary Information

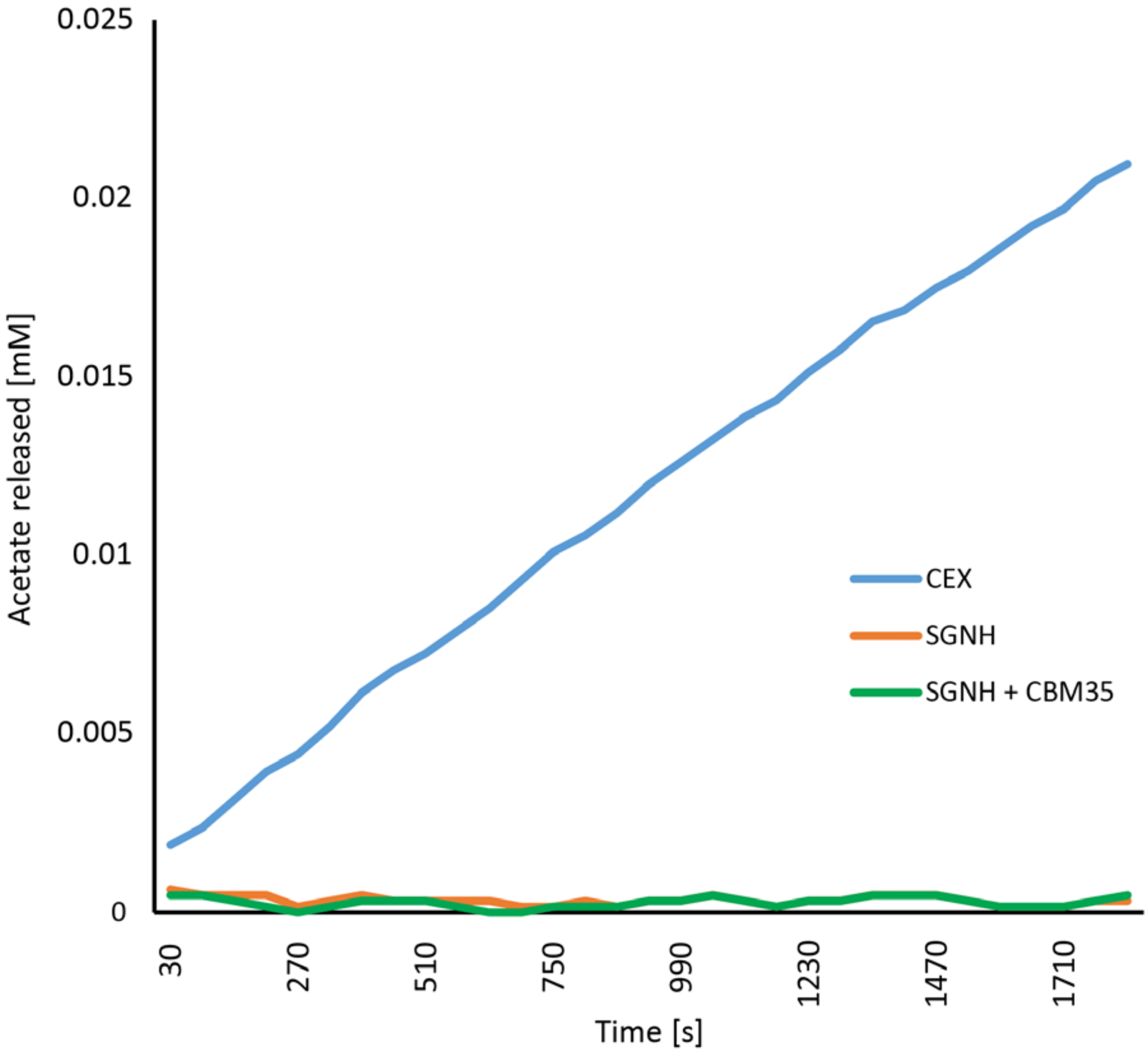


**Fig. S1** Activities of the SGNH domain (orange trace) and a cocktail of SGNH domain with the CBM35 (green trace) as two separate peptides were tested on *p*NP acetate, in comparison with a complete *Ri*CEX (blue trace). The conditions (pH 7.0 and 30 °C) were selected based on the pH optimum recorded previously. Despite using a much higher protein concentration in the single-domain samples (403 nM of the SGNH domain, 606 nM of the CBM35, as compared to 100 nM of *Ri*CEX), no activity was observed.

**Protein thermal shift assay**

*Ri*CEX showed great stability and negligible loss of activity after long term storage at 4°C, pH 8.0, while *Ri*CE2 was relativly unstable and required storage at pH 5.9, -80 °C. To investigate the optimum storage pH and thermal stability of the esterases, we examined the denaturation of *Ri*CEX and CE2 in buffers at pH 5.0-8.0 using the Protein thermal shift (PTS) assay (Thermo Scientific, USA). *Ri*CEX was stable up to 65 °C, with the highest observed melting temperature at pH 8.0. *Ri*CE2 has a markedly lower stability when exposed to increased temperatures, with a melting point of 39 °C at pH 8.0 and a highest melting point of just 47°C at pH 6.0. The CBM35 domain does not fold properly on its own (Fig. S2C), while the SGNH domain appears to fold into a slightly less stable structure than the complete enzyme (denaturing at 52 °C at pH 8.0, as compared with 65 °C for the complete *Ri*CEX). This indicates that both domains may be necessary for correct folding and activity.

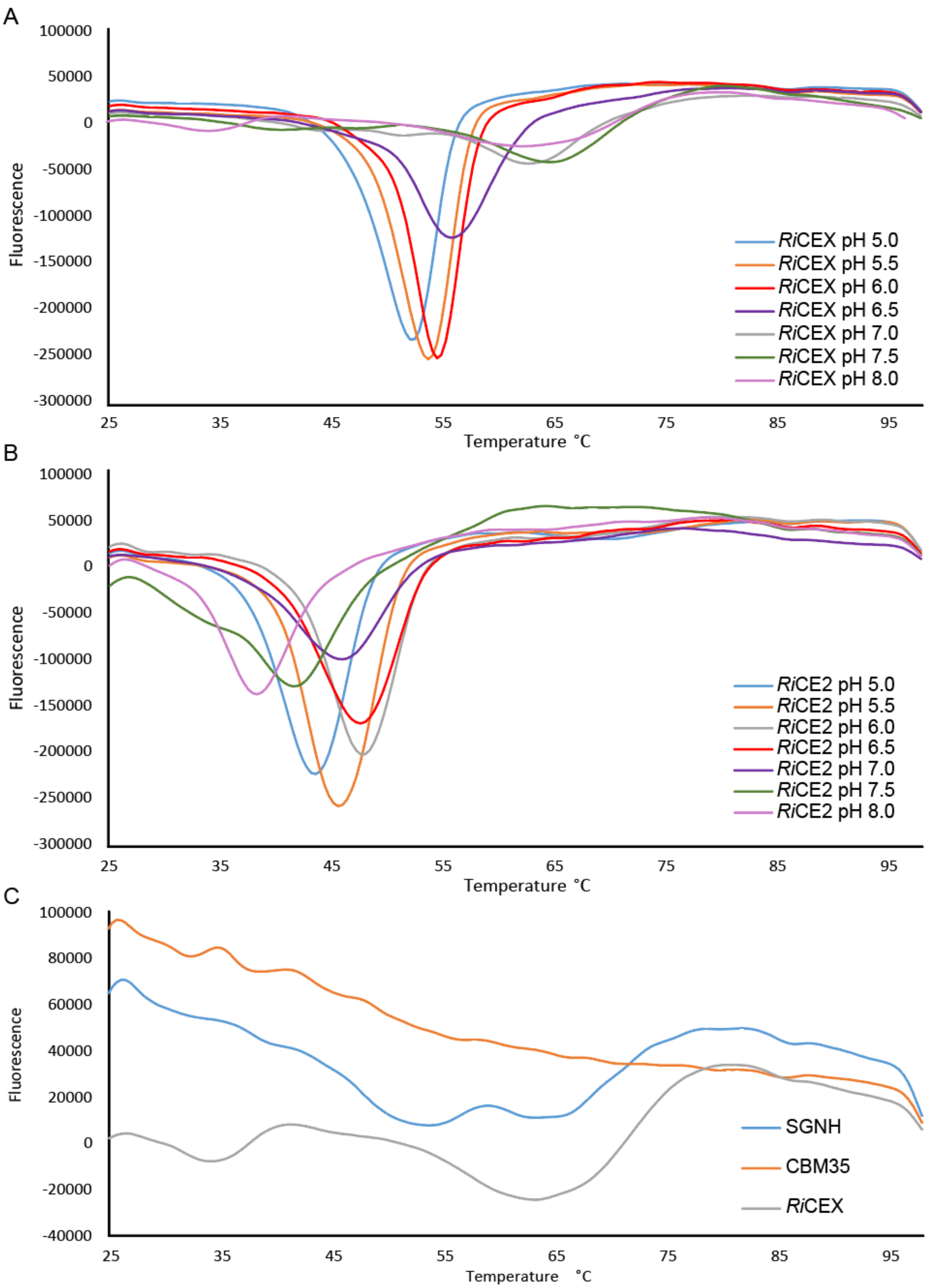


**Fig. S2.** Melt plots of derivative data from protein thermal shift assays. Lowest point in each plot signifies the temperature point at which the protein is denatured. (A) The highest melting temperature (64 °C) for *Ri*CEX was observed at pH 7.5. (B) The highest melting temperature (48 °C) for *Ri*CE2 was observed at pH range 6.0- 6.5. (C) The SGNH domain appears to fold into a stable complex, with its melting temperature at 53 °C, while the CBM35 appears to hold no particular structure (melting temperature measured at pH 8.0).

**HMM building**

In order to build Hidden Markow Models (HMMs), the phmmer tool (1) on the European Bioinformatics Institute website (www.ebi.ac.uk) was used to search for homologous sequences in the UniProt's reference proteomes database. Homologous sequences were used to generate an HMM (2) which was then used to search the UniProtKB (3) database for more distant relatives of the *Ri*CEX.

Searching the UniProtKB (3) using the database with the CBM35 domain of *Ri*CEX using a protein vs protein database tool phmmer (with significance threshold of $e^{-20}$) resulted in 154 sequences of which 135 came from Firmicutes and were found in the same two domain arrangement with a Lipase_GDSL_2 domain. These results indicate that the CBM35 structure might be specific to the phylum.

A phmmer search in the UniProtKB reference proteomes using the whole sequence of *Ri*CEX with an E-value for significant hit at $1e^{-99}$ resulted in 32 sequences, all of which had the same two domain architecture and originated from Firmicutes. These were used for HMM building. A search in the UniProtKB database with an E-value $<1e^{-50}$ using this HMM returned 482 bacterial sequences with a two domain GDSL2_Lipase + CBM35 architecture. The majority of these sequences came from Firmicutes (435). 16 results came from Bacteroidetes. The sequences of those 482 sequences were aligned to identify conserved aminoacids. All three catalytic residues (serine 41, aspartic acid 190, histidine 193) and the tryptophan 326 of the clamping domain that orients the substrate in the active site by aromatic stacking were present in the 90% consensus sequence (Fig. S3). The presence of these residues in consensus sequences indicates that they are conserved, and could be considered characteristic for the esterase family. Aligned sequences of five esterases homologous to *Ri*CEX from known polysaccharide degraders: A0A1R1F575 of *Paenibacillus rhizosphaerae,* A0A1G5RU69 of *Pseudobutyrivibrio xylanivorans,* D4KAW7 *Faecalibacterium prausnitzii* SL3/3, A0A2S6HWM2 *Bacteroides xylanolyticus,* D9SWK7 *Clostridium cellulovorans* (protein names are Uniprot accession numbers) (Fig. S4) show that amino acid residues important for activity are conserved among members of diverse species, even in enzymes sharing low identity.

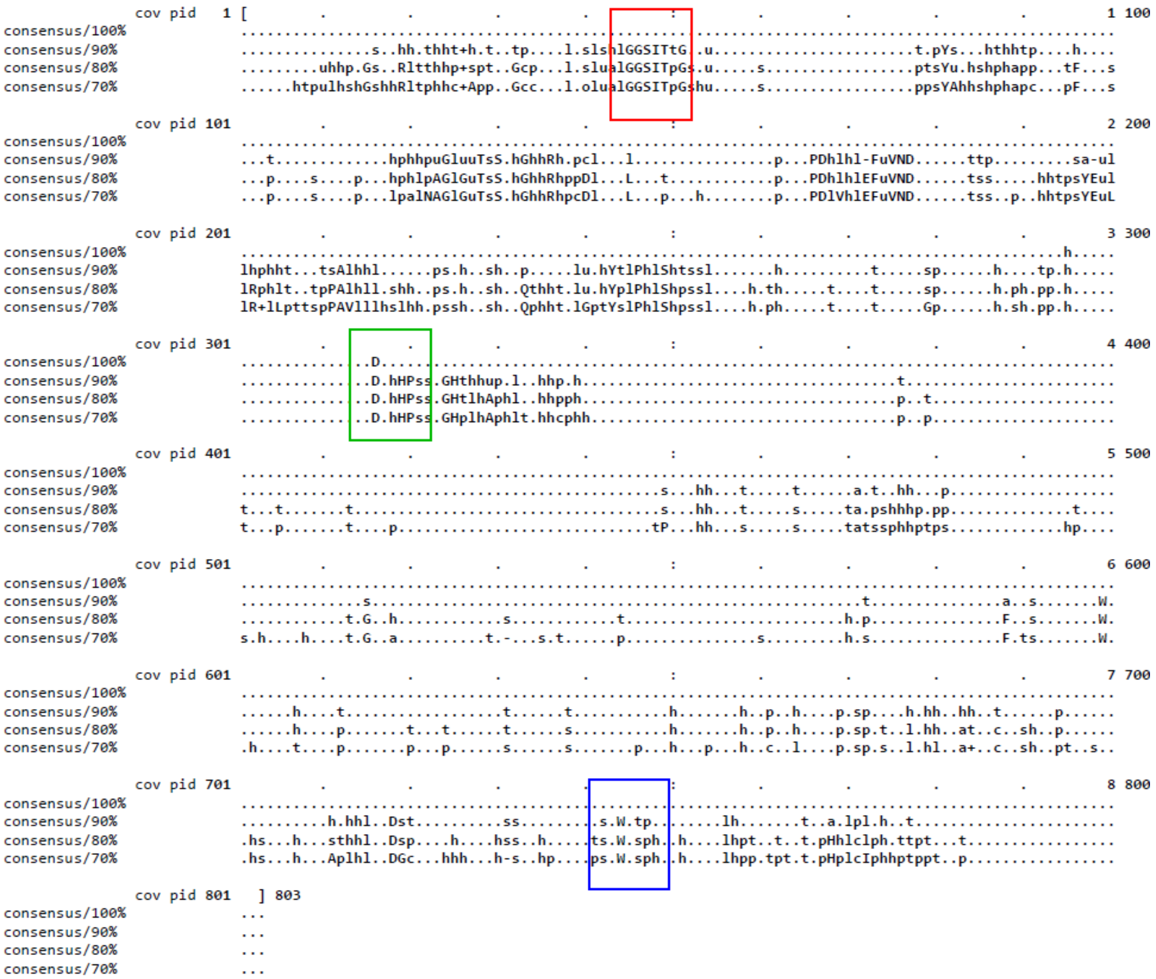


**Fig. S3.** MView generated consensus sequences of the 482 sequences found in UniprotKb database using an HMM of the *Ri*CEX. Homologues of all three catalytic residues of *Ri*CEX (Ser41 in the red box, Asp190 and His193 in the green box) as well as the Trp326 of the CBM35 domain (blue box) are present in the 90% consensus sequences. Capital letters signify conserved aminoacids, small letters indicate conserved amino acid characters as described by Taylor et al. (4): l – aliphatic, a – aromatic, c – charged, h – hydrophobic, - – negative, p – polar, + – positive, s – small, u – tiny, t – turnlinke.

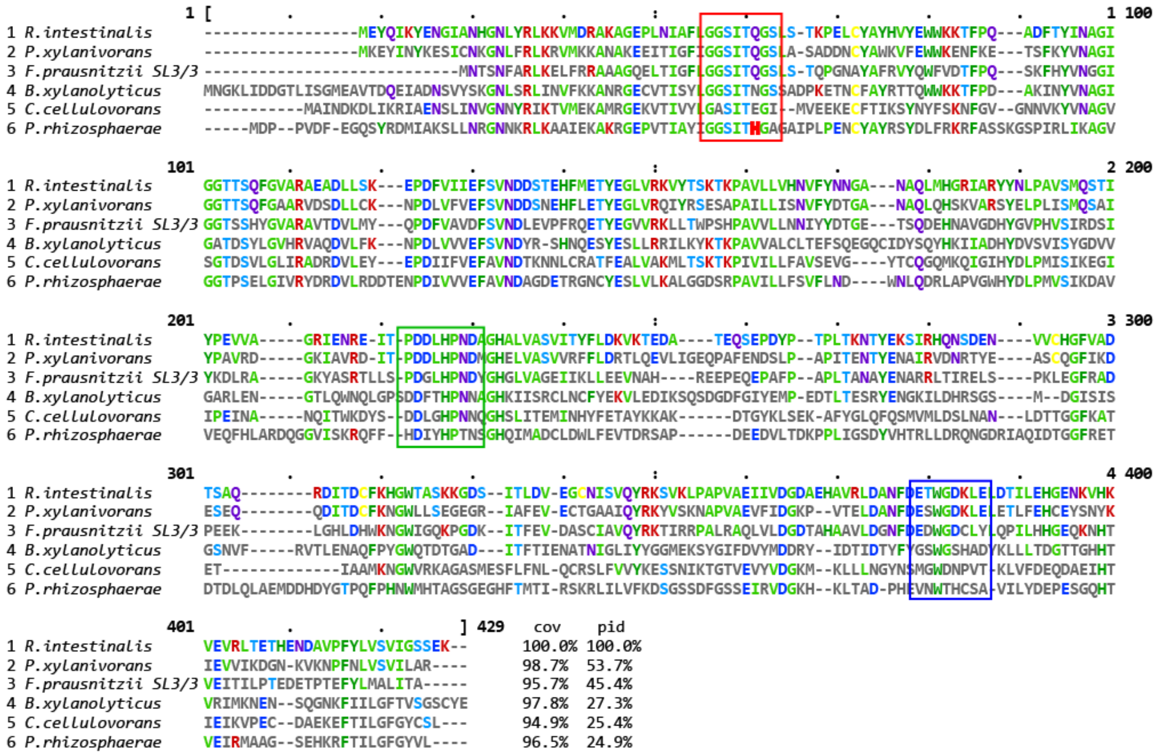


**Fig. S4**. MView generated alignment of six *Ri*CEX homologues from six distinct taxonomic units (1, *Roseburia intestinalis*; 2, *Pseudobutyrivibrio xylanivorans*; 3, *Faecalibacterium prausnitzii* SL3/3; 4, *Bacteroides xylanolyticus; 5, Clostridium cellulovorans;* 6, *Paenibacillus rhizosphaerae*) found in the UniProtKB HMM search. A closer look at the *Ri*CEX homologue found in *Bacteroides xylanolyticus* (Uniprot A0A2S6HWM2) showed that this protein is associated with a cluster for β-mannan degradation, including an ABC-transporter typical for Firmicutes. This organism has no Sus-system, that is typical for Bacteroides, indicating that it has been incorrectly annotated and rather part of the Firmicutes phylum. Residues are colored with the Clustal color scheme (5) based on conserved amino acid chemical character. Cov – coverage, pid – protein identity compared to *Ri*CEX.

Table S1. Crystal data, data collection, and refinement statistics.

| | | |
|---|---|---|
| *Crystal data* | | |
| | *Ri*CEX | *Ri*CEX-mannopentaose |
| Space group | P 1 $2_1$ 1 | |
| Crystal parameters | a = 75.12, b = 135.52, c = 85.13 | a = 75.48, b = 136.69, c = 85.41 |
| | $\alpha$ = 90, $\beta$ = 115.09, $\gamma$ = 90 | $\alpha$ = 90, $\beta$ = 113.86, $\gamma$ = 90 |
| *Data collection* | | |
| X-ray source | ESRF, ID23-1 | ESRF, ID23-1 |
| Resolution (Å)[a] | 48.2-1.75 (1.78-1.75) | 48.6-2.40 (2.53-2.40) |
| Wavelength (Å) | 0.97531 | 0.97625 |
| Temperature (K) | 100 | 100 |
| Number of unique reflections | 155949 (7703) | 61043 (8942) |
| Completeness[a] | 100 (99.9) | 98.7 (99.2) |
| Redundancy[a] | 6.7 (6.4) | 3.5 (3.5) |
| CC half[a] | 0.998 (0.815) | 0.995 (0.713) |
| I/ $\sigma$(I)[a] | 11.5 (2.2) | 8.2 (2.0) |
| $R_{sym}$[b] | 0.079 (0.631) | 0.097 (0.593) |
| *Refinement statistics* | | |
| $R_{cryst}$[c] | 0.190 | 0.181 |
| $R_{free}$[d] | 0.232 | 0.253 |
| Wilson B-factor (Å$^2$) | 25.4 | 42.1 |
| Ramachandran plot,in most favored/other allowed regions (%) | 96/4 | 95/5 |
| Standard Uncertainty (Maximum Likelyhood): | 0.093 | 0.224 |
| Added waters | 808 | 487 |
| **PDB code** | 6HFZ | 6HH9 |

[a] *Values for outer shell in parenthesis,*

[b] $R_{sym} = \Sigma|I - \langle I\rangle|/\Sigma I$.

[c] $R_{cryst} = \Sigma(|F_{obs}| - |F_{calc}|)/\Sigma|F_{obs.}|$

[d] *$R_{free}$ is the $R_{cryst}$ value calculated on the 5 % reflections excluded for refinement.*

**Esterase activity**

Substrate specificity of the two esterases was tested on a wide range of relevant substrates. Activity was only observed on mannose-based oligosaccharides, and the patterns of activity were always similar: a partial deacetylation when either esterase was used, and a near complete deacetylation when both enzymes were used (Fig. S5 A-C, and Fig. 4). None of the esterases were active on cellulose monoacetate, acetylated xylan and chitin oligosaccharides (Fig. S5 D-F).

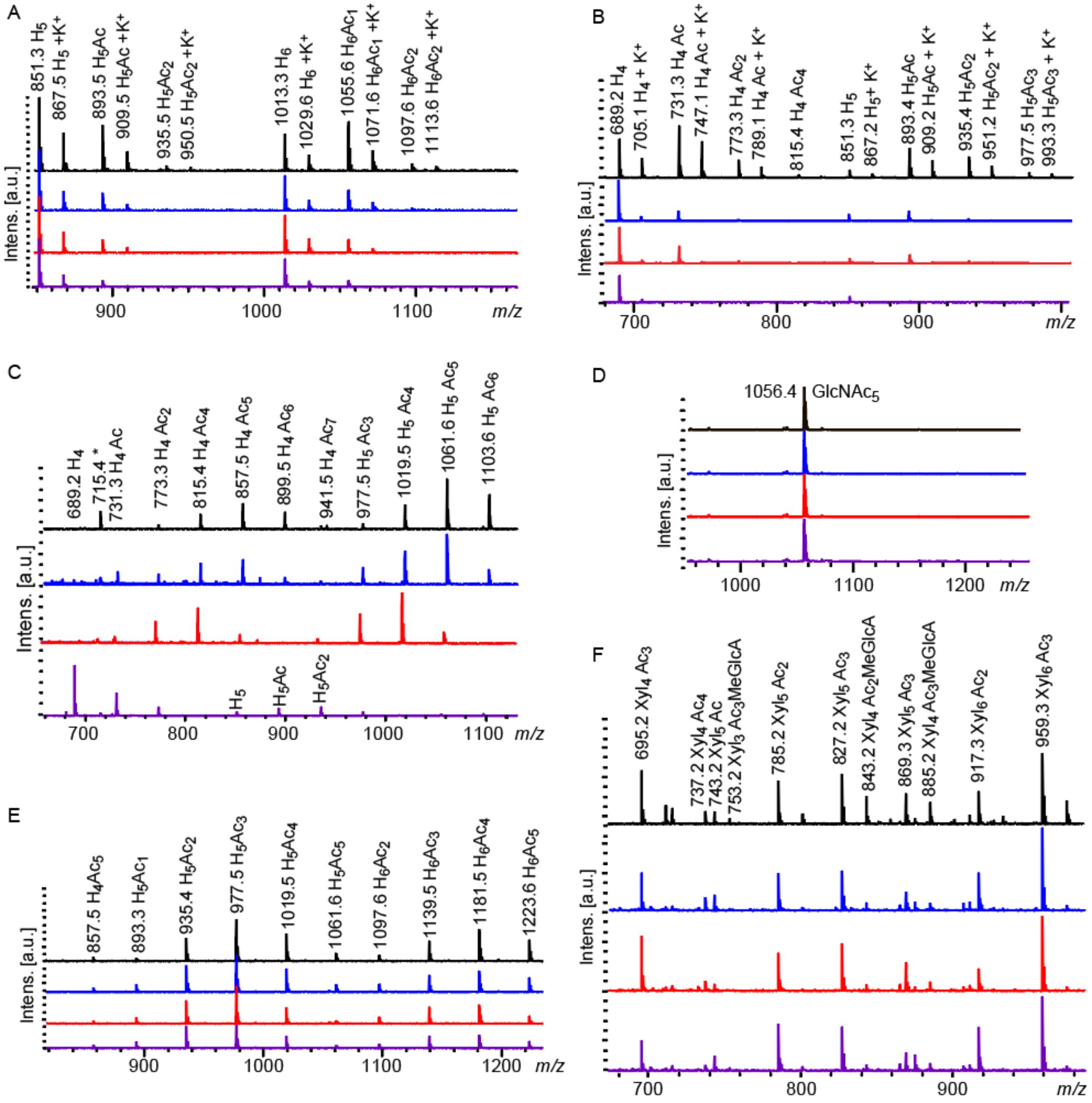


**Fig. S5**. MALDI ToF traces of enzyme reactions on various substrates. (A) Konjac glucomannan deacetylated with *Ri*CE2 (in blue), *Ri*CEX (in red), and both enzymes combined (in purple). (B) Chemically acetylated Konjac glucomannan deacetylated with *Ri*CE2 (in blue), *Ri*CEX (in red) and both enzymes combined (in purple). (C) Aloe vera mannan treated with *Ri*CE2 (in blue), *Ri*CEX (in red) and the two enzymes in combination (purple). (D) Chitopentaose (penta-N-acetylchitopentaose) (Megazyme, Ireland) treated with *Ri*CE2 (in blue), *Ri*CEX (in red) and both enzymes combined (in purple), showing no signs of activity. (E) Acetylated cellulose oligosaccharides treated with *Ri*CE2 (in blue), *Ri*CEX (in red) and both enzymes combined (in purple), the spectra show no signs of enzymatic activity. (F) Birch xylan oligosaccharides treated with *Ri*CE2 (in blue), *Ri*CEX (in red) and both enzymes combined (in purple) showing no apparent activity on the xylooligosaccharides H-hexose, X-xylose, Ac-acetylations, Me-methylations, GlcA-glucuronic acid, $GlcNAc_5$ - Chitopentaose. All masses represent sodium adducts unless marked with K+, unlabeled peaks represent background signals from the sample matrix. Peak labelled 715.4* in panel C is a persistent contaminant.

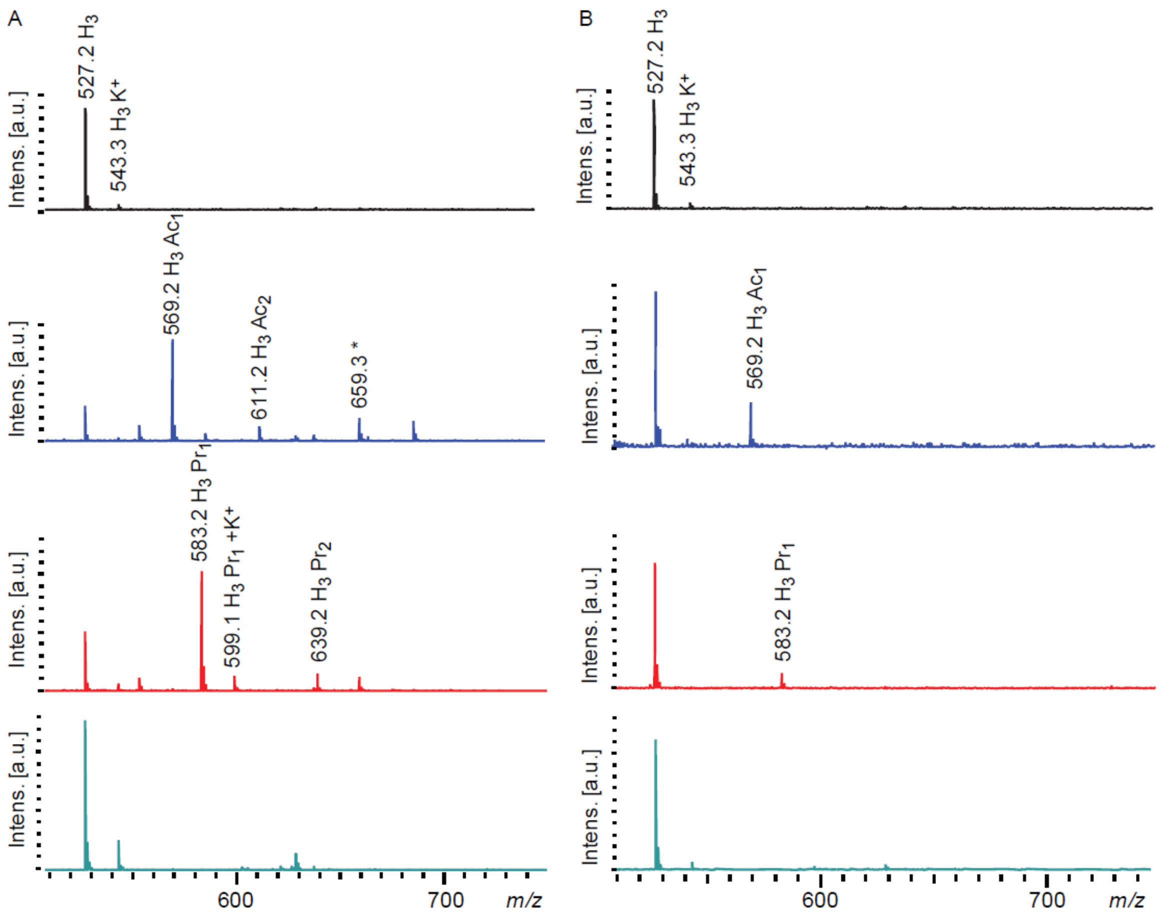


**Fig. S6.** MALDI-ToF spectra of transesterification reactions with mannotriose and vinyl esters. (A) *Ri*CE2 was able to transacetylate (in blue) and transpropylate (in red) mannotriose (in black), but not transbutyrylate (in green). (B) Similarly, *Ri*CEX was able to transacetylate (in blue) and transpropylate (in red) mannotriose, but was not able to transbutyrylate it (in green). Abbreviations: H- hexose, Ac- acetylation (*m/z* 42), Pr- propylation (*m/z* 56), $K^+$ signifies peaks of potassium adducts, all other *m/z* are of sodium adducts. Peak labelled * in (A) (*m/z* 659.3) is a persistent contaminant.

### Acetyl migration:

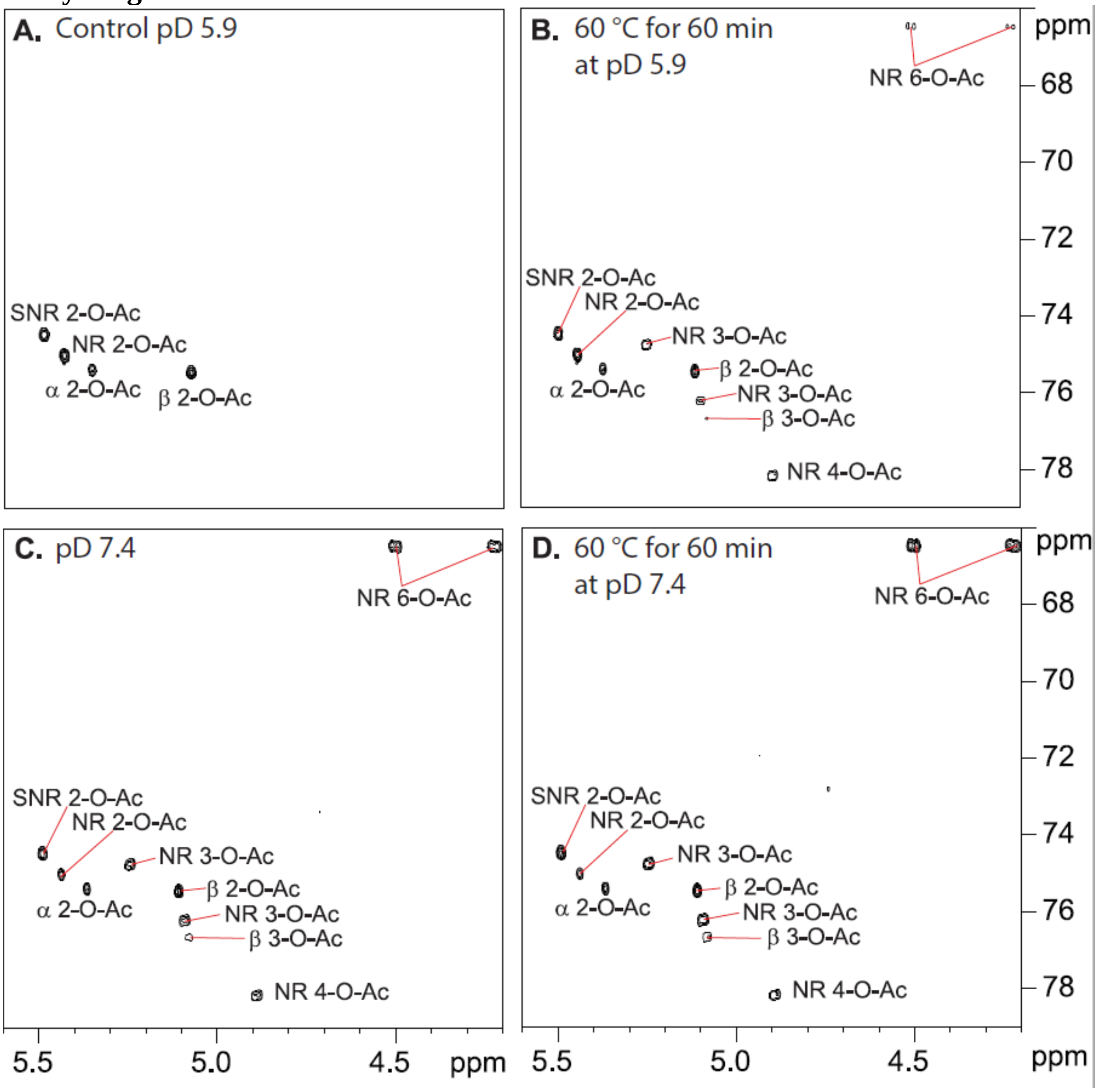


**Fig. S7.** Acetyl migration on *Ri*CEX transacetylated oligosaccharides (A) n 20 mM sodium phosphate *Ri*CEX at pD 5.9 acetylated mannotriose exclusively on the hydroxyl groups on carbon C2, (B) after exposure to 60°C for one hour at pD 5.9, (C) after pD was increased to 7.4 by adding sodium phosphate and (D) after exposure to both high temperature and pD 7.4. In all conditions, the acetyl groups, initially present only as 2-*O*- acetylations migrated to other sites on the same sugar units. NR - nonreducing end mannose, SNR – Mannose internal in the mannotriose, α and β – anomeric configuration of the reducing end mannose.

The experiments with *Ri*CEX transacetylated mannotriose have shown that temperatures of just 60 °C and pD 7.4 induce acetyl migration. On the 2-*O*- acetylated mannotriose, the apparent direction of migration was the 'clockwise' 2-*O*- → 3-*O*-. To test if acetyl migration is a directional process, we devised a set of experiments complementary to those described in the NMR analysis described in Fig. S5. A sample of GGM was treated overnight with excess *Ri*CEX (1 µM final concentration in sample) to remove all 2-*O*-acetylations, leaving a substantial amount of acetylation and exposing non-acetylated 2-*O*- hydroxyls as destination for 3-*O*- → 2-*O*- migrating acetylations. The release of acetate forced the pH of the solution to 5.6, ensuring that no unwanted migration was occurring during the following handling of samples. *Ri*CEX was then removed from the solution by filtration, and the solution was split into two samples: one for pH induced migration, and one for temperature induced migration. pH induced migration was accomplished by adjusting the sample pH to 7.4 by slow addition of a sodium phosphate buffer at pH 7.75, and incubation at 30 °C overnight. MS analysis was carried out at every step to ensure that deacetylation was the result of 2-*O*- acetylations being enzymatically removed rather than chemical deacetylation. Temperature induced migration was accomplished by adjusting the pH to 5.9, and incubation at 60 °C for one hour. Conditions in both experiments were selected to match those in the 2-*O*-→ 3-*O*- migration experiment (Fig. S5) described above. After the migration was induced, samples were treated with 1 µM final concentration of *Ri*CEX again, saving a portion of the original sample as control, before analysis by MALDI ToF MS. The cycle of enzymatic treatment, enzyme removal, pH adjustment, migration induction, enzymatic treatment and analysis was completed twice for the pH induced migration and three times for the temperature induced migration before the acetylations were completely removed from the mannan. Both pH (Fig. S6) and heating (Fig. S7) induce a migration process which replaced the 2-*O*- acetylations, providing fresh substrates for deacetylation by *Ri*CEX at each round, until a complete deacetylation was achieved. Reaction rates for the acetyl migration between each pair of adjacent acetylations were presented in literature (6).

The rate of 2-*O*- →3-*O*- migration as well as the reverse rate were the highest (0.566 $h^{-1}$ and 0.395 $h^{-1}$ respectively at pD 8.0, 25℃) of all migration steps (6). Inducing migration on a mixture of heterogeneous mannooligosaccharides with 2-*O*- acetylations selectively removed allowed us to demonstrate that the migration occurs in all directions depending on the distribution of remaining acetylations. The elevated temperature and pH do not drive this reaction in a particular direction, but rather create a permissive environment where the acetylations can re-distribute to their most energetically favourable conformation. The apparent direction of acetyl migration depends on the distribution of acetylations prior to exposure to migration permissive conditions. Migration experiments presented here show that this process occurs both in model substrates and heterogeneous mixtures of hemicellulose obtained from a typical industrial hydrothermal extraction.

These results show that acetylations readily redistribute to an equilibrium when exposed to conditions that facilitate migration, such as heating or pH>6.0. This finding is especially important for hemicellulose biorefining with enzymatic deacetylation steps – since conditions throughout the process can quickly change the acetyl distribution. As acetylations affect the solubility and viscosity of mannans in solution, selective enzymatic deacetylation and easily achieved redistribution could be used for fine-tuning the physicochemical properties of mannans for hydrocolloid applications. This observation has a special significance for biorefining of mannan rich feedstocks such as softwood mannans. Migration induced by a short exposure to just 60 °C, at pH 5.9 as well as previously published data on migration caused by heating (7) imply that the distribution of acetylations present in hemicellulose produced by steam explosion and other common hydrothermal extraction methods may not represent the distribution of acetylations present in the hemicellulose *in vivo.*

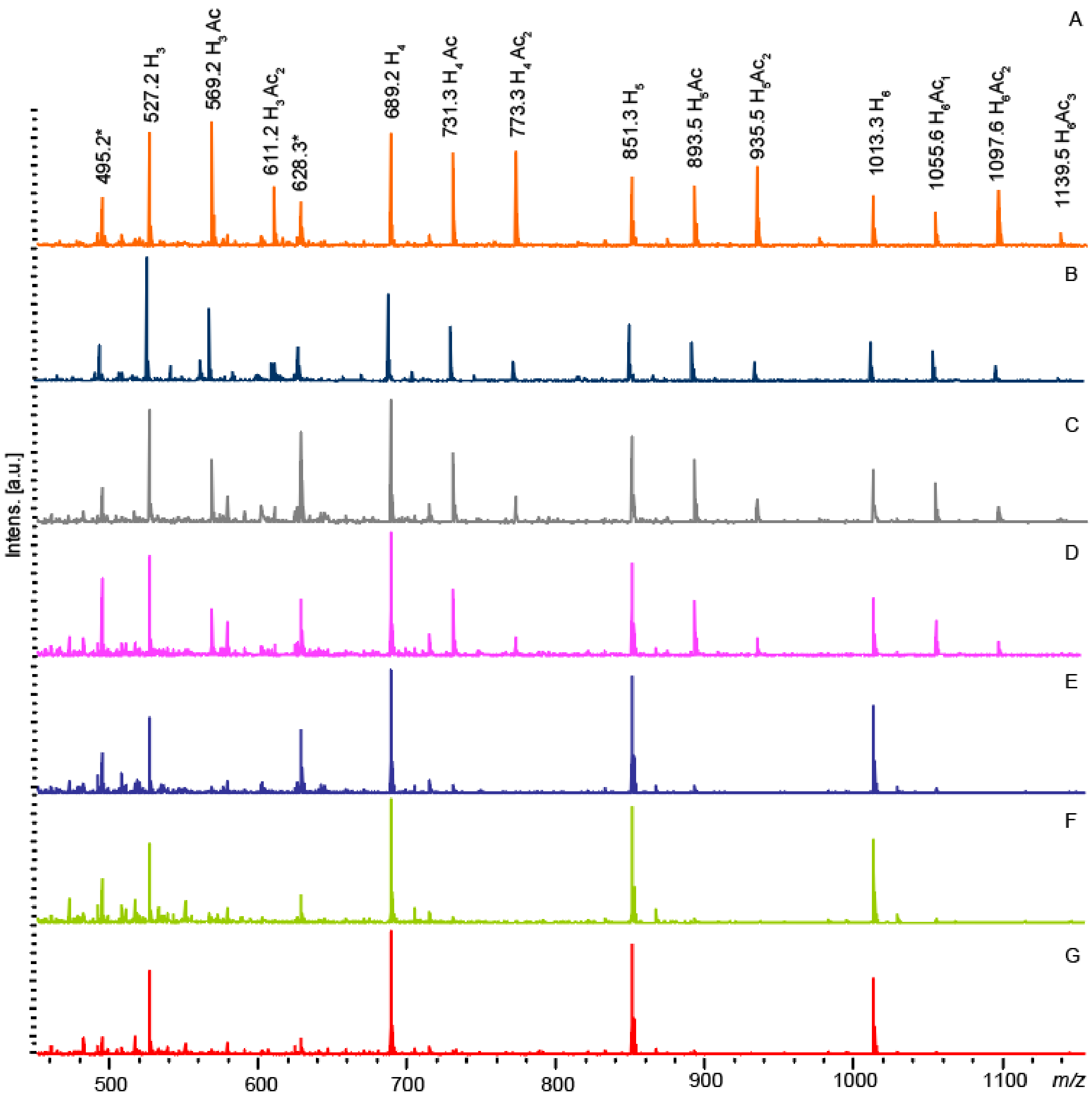


**Fig. S8.** MALDI ToF MS spectra of Norway spruce GGM at each stage of pH induced migration. (A) The original sample had a prevalence of multiple acetylated oligosaccharides. (B) Overnight treatment with *Ri*CEX removed a significant portion of the acetylations. (C) After *Ri*CEX was removed by filtration, pH of the sample was adjusted to 7.4, there was no apparent deacetylation in the process. (D) The sample retained the acetylations after incubation at 30 °C overnight. (E) Treating the overnight incubated sample with *Ri*CEX resulted in removal of a further portion of acetylations, with minor peaks for single acetylated oligosaccharides present. (F) Repeating the treatment in (C) did not affect the deacetylation. (G): Treating the sample in (F) with *Ri*CEX for the third time resulted in complete deacetylation.

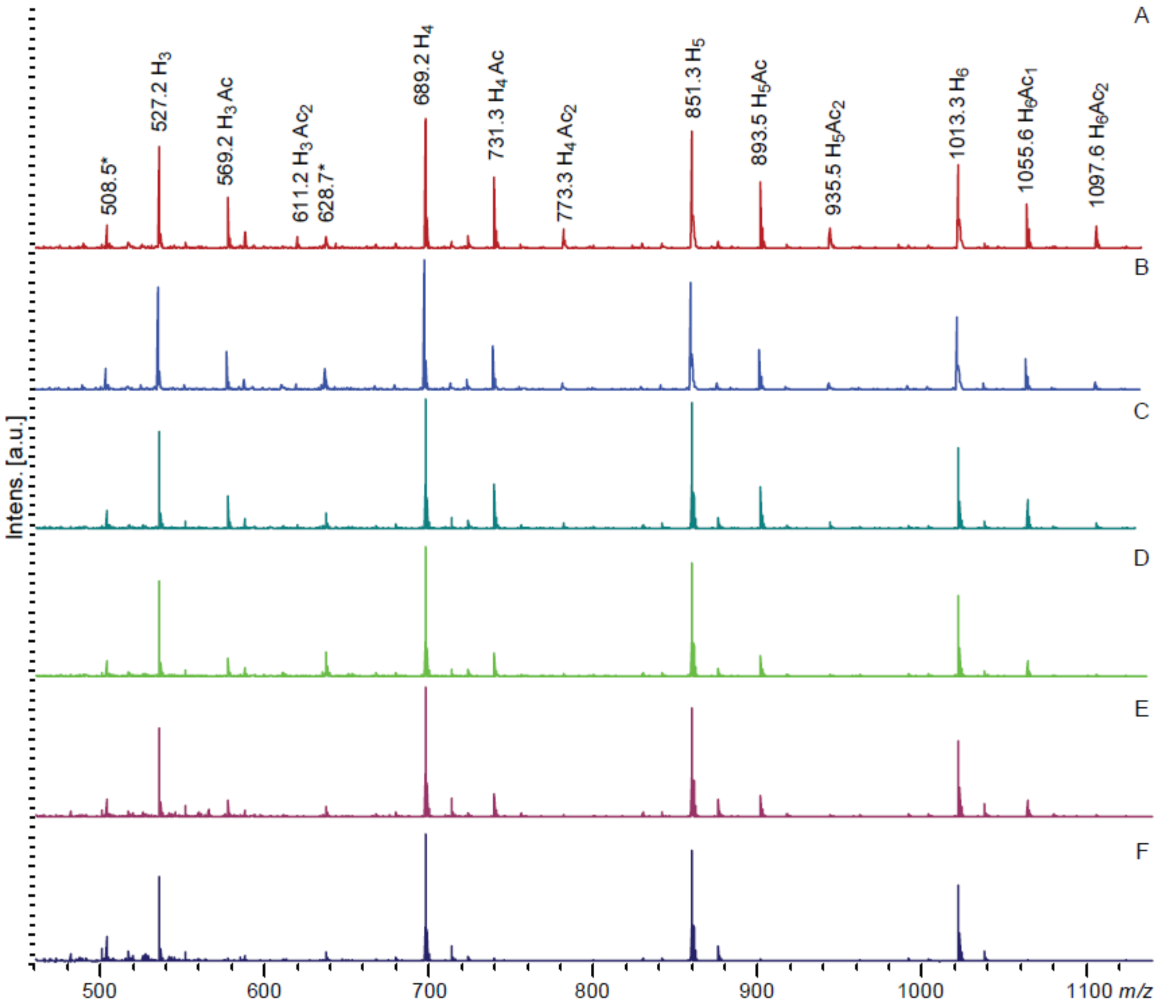


**Fig. S9.** MALDI ToF spectra of Norway spruce GGM at each step of temperature induced migration. (A) After the initial deacetylation and filtration, the pH of the sample adjusted to 5.9, and the sample was incubated at 60 °C for one hour without a loss of acetylations in the process. (B) *Ri*CEX treatment of the heat treated sample removed more acetylations. (C) After *Ri*CEX was removed by filtration, the sample pH was adjusted to 5.9, and the sample was incubated at 60°C for the second time, with no loss of acetylations. (D) After the second heat treatment, *Ri*CEX removed a further proportion of acetylations, with peaks for double acetylated oligosaccharides disappearing completely. (E) The control sample after the third round of heat-induced migration (as in (C)). (F) After three rounds of heat-induced migration, the *Ri*CEX treatment removed all acetylations.

**SI Materials and Methods:**

1. Cloning and expression of *Ri*CEX truncated versions.

*Ri*CEX gene fragments were amplified from *R. intestinalis* genomic DNA using the primer pairs CEX_up/CEXcatD_rev1 and CEXcbm_up/CEX_down, respectively (Table S2). Fragments were cloned into the pNIC-CH expression vector with a C-terminal hexa-histidine tag by ligation-independent cloning (LIC) (8), giving the constructs pNIC-*Ri*CEX$_{CATD}$ and pNIC-*Ri*CEX$_{CBM35}$. Transformants were verified by sequencing. The two proteins were expressed in *Escherichia coli* BL21(DE3) cells harboring the appropriate recombinant plasmids. The recombinant *E. coli* strains were pre-cultured overnight in Luria Bertani (LB) broth supplemented with 50 µg/mL kanamycin (Sigma-Aldrich, Germany) and then used to inoculate 500 mL of medium consisting of 450 mL LB, 50 µg/mL kanamycin and 50 mL of potassium phosphate buffer (0.17 M $KH_2PO_4$, 0.72 M $K_2HPO_4$). Protein expression was induced by adding isopropyl-β-D-thiogalactopyranoside (IPTG) to a final concentration of 0.5 mM for 16 h at 23°C. Cells were harvested by centrifugation (6000 g for 10 minutes) and resuspended in 30 mL lysis buffer (50 mM Tris-HCl, pH 8.0, 500 mM NaCl, 10 mM imidazole). A cell-free extract was prepared by pulsed sonication and centrifugation at 15000 g for 15 minutes. The supernatant containing the soluble proteins was collected and filtered with 0.22-µm syringe filters. Recombinant *Ri*CE2 and *Ri*CEX were purified by immobilized metal affinity chromatography (IMAC) and size exclusion chromatography (SEC). Protein purity was verified by sodium dodecyl sulfate-polyacrylamide gel electrophoresis (SDS-PAGE). Protein concentration was determined using the Bradford assay (Bio-Rad) with bovine serum albumin as a standard.

**Table S2: Primers used in this study. 5' extension sequences used for molecular cloning are underlined.**

| Gene | Primer (5' -3') |
|---|---|
| CEX_up | **F:**TTAAGAAGGAGATATACTATGGAATATCAAATTAAATACGAAAACGGC |
| CEXcatD_rev1 | **R:**AATGGTGGTGATGATGGTGCGCCTGTTCCGTTGCATCTTCTG |
| CEXcbm_up | **F:**TTAAGAAGGAGATATACTATGGATTATCCGGCACCTCTCAC |
| CE2_down | **R:**AATGGTGGTGATGATGGTGCGCAGATTCCCAGACTGCATCCC |

2. *Ri*CEX crystallography

For crystallographic studies, seleno-L-methionine substituted *Ri*CEX was produced by first transforming pNIC-*Ri*CEX (9) into *E. coli* B834(DE3) cells via the heat-shock method and plating the bacteria onto LB agar supplemented with 50 µg/mL kanamycin. Recombinant cells were grown in Medium A according to the EMBL protocol (https://www.embl.de/pepcore/pepcore_services/protein_expression/ecoli/seleno/), supplemented with 50 µg/mL methionine (Sigma-Aldrich, Germany) and 50 µg/mL kanamycin at 37°C until an $OD_{600}$ of 0.8 was reached. At this point, cells were harvested by centrifugation at 6000 g for 15 minutes and resuspended in an equal amount of kanamycin-supplemented Medium A. Temperature was adjusted at 23 °C and, following a 2 h starvation period, the flask was supplemented with 50 µg/mL seleno-L-methionine (Sigma-Aldrich, Germany). After 30 min of further incubation, seleno-L-methionine substituted *Ri*CEX expression was induced with 0.5 mM IPTG and cultures were allowed to grow for an additional 48 h before being collected. Cells were harvested by centrifugation at 5000 g for 10 minutes, cell pellets were resuspended in 30 mL of 50 mM Tris buffer containing 10 mM Imidazole and 500 mM NaCl and lysed by sonication. The cell lysate was centrifuged at 12000 g for 15 minutes and the supernatant containing the proteins was collected, sterile filtered and purified as described above.

3. Preparative HPLC.

Preparative chromatography was conducted using an Agilent 1260 Infinity preparative chromatography system with an XBridge BEH prep OBD 5 µm particle size 30x250mm column. Analytical HILIC method was scaled up to 17.5 mL/min flow with 3.85 mL injections of 1-5 mg/mL carbohydrate concentrations. Elution started with a step of 0 – 3.57 min 75 % acetonitrile, followed by a linear gradient of 3.58 – 14.28 min 75 % - 50 % acetonitrile, a step of 14.29 - 21.42 min 50 % acetonitrile, and a final step of 21.43 – 33 minutes at 75 % acetonitrile. Fractions were collected as one minute time slices, acetonitrile evaporated in a fume hood and liquid fractions freeze-dried.

4. Transesterification.

Transesterification of oligosaccharides was conducted using vinyl acetate, vinyl butyrate and vinyl propionate (Thermo scientific, USA) as acyl donors. Enzymes were added to oligosaccharide solutions with concentrations from 1 – 10 mg/mL, and a volume of vinyl esters equal to 20 – 50 % sample volume was added. The samples were incubated in a thermomixer (Eppendorf, Norway) shaking at 600 rpm overnight, then moved to a freezer at -20 °C. The vinyl acetate, which remained liquid on top of the frozen aqueous phase was removed from the samples, and 96% ethanol was added on top of the frozen sample until the final concentration exceeded 80% in order to deactivate the enzymes. Samples were then thawed by vortexing and filtered through a pre-washed 1mL Amicon Ultracel 3kDa ultrafiltration device (Merck KGaA, Germany) to remove the enzymes completely and minimize the risk of deacetylation. Oligos were then dried in an Eppendorf Concentrator plus (Eppendorf, Norway) at room temperature.

For the purpose of preparative chromatography, samples were frozen, liquid layer of vinyl acetate was removed and samples were then diluted with acetonitrile to a 75 % concentration, foregoing the filtration step.

5. NMR

To reduce the interference of the water signal the substrate, *Ri*GH26 treated spruce galactoglucomannan was dissolved in 99.9% $D_2O$ (Sigma-Aldrich, Germany) and lyophilized. Similarly, 10 mL 40 mM phosphate buffer pH 5.9 and 250 mM phosphate buffer pH 8.0 were lyophilized and the powder were dissolved in 10 mL 99.9% $D_2O$.

For the time-resolved NMR recordings: 4-5 mg of Norway spruce galactoglucomannan hydrolyzed with the *Ri*GH26 mannanase or *Ri*CEX transacetylated mannotriose were dissolved in 500 µL 40 mM phosphate buffer pD 5.9 (99.9% $D_2O$) and transferred to a 5 mm NMR tube. The sample was preheated in the NMR spectrometer for ~10 min. Hereafter all recording parameters were set prior to the time-resolved

NMR experiment. 2 or 5 µL of enzyme solution (to a final concentration of 1 µM *Ri*CEX or 10µM *Ri*CE2) was added to the preheated substrate and mixed by inverting the sample three times. The sample was then immediately inserted into the preheated NMR spectrometer and the experiment was started (time from adding the enzyme to the first spectra has been recorded was between 3-4 minutes in total). The recorded spectrum is a pseudo-2D type experiment recording a 1D proton NMR spectrum with weak water suppression (Based on Bruker 1D proton setup for metabolomics noesygppr1d) every 5 min with in total 200 time points. The recorded 1D proton spectrum contains 32K data points and has a spectral width of 10 ppm, 24 scans, and pre-saturation during relaxation delay and 10 ms mixing time with spoil gradient and relaxation delay of 1 s (total recording time of 89s).

To monitor the effect of temperature and pH on acetyl migration, 2 mg transacetylated mannotriose was dissolved in 500 µL 40 mM phosphate buffer pD 5.9 (99.9% D2O) and it served as a control sample where 1D proton and 2D $^{13}C$ heteronuclear single quantum coherence (HSQC) with multiplicity editing spectra were recorded. The sample was split into 3 samples of 160 µL each and transferred into 3mm NMR tubes. Hereafter, the first sample was heated to 60 °C for 60 min. In the second sample pD was adjusted pD 7.4 by adding 20 µL of 250 mM phosphate buffer pD 8.0 and in the third sample the pD was adjusted pD 7.4 by adding 20 µL of 250 mM phosphate buffer pD 8.0 and heated to 60 °C for 60 min. A 1D proton and 2D $^{13}C$ HSQC spectra were recorded at 25 °C for each of the samples.

All homo and heteronuclear NMR experiments were recorded on a BRUKER AVIIIHD 800 MHz (Bruker BioSpin AG, Switzerland) equipped with a 5 mm cryogenic CP-TCI. All NMR recordings were performed at 37°C. For chemical shift assignment of *Ri*CEX transacetylated mannotriose, the following spectra were recorded: 1D proton, 1D proton with presaturation during relaxation delay and 10 ms mixing time with spoil gradient, 2D double quantum filtered correlation spectroscopy (DQF-COSY), 2D total correlation spectroscopy (TOCSY) with 70 ms mixing time, 2D $^{13}C$ HSQC, 2D $^{13}C$ Heteronuclear 2 Bond Correlation (H2BC), 2D $^{13}C$ HSQC-[$^{1}H$,$^{1}H$]TOCSY with 70 ms mixing time on protons and 2D heteronuclear multiple bond correlation (HMBC) with BIRD filter to suppress first order correlations. The water signal to 4.75 ppm (at 25 °C, pH 5.5 (10)) was used as chemical shift reference for protons, while $^{13}C$ chemical shifts were referenced indirectly, based on the absolute frequency ratios (11). The spectra were recorded, processed and analyzed using TopSpin 3.5 software (Bruker BioSpin AG, Switzerland).
